# How variable and stress-sensitive is sleep expression in lizards?

**DOI:** 10.64898/2026.09.01.748758

**Authors:** Nitya Prakash Mohanty, Paul-Antoine Libourel, Dhanya Bharath, Mihir Joshi, Maria Thaker

**Affiliations:** Centre for Ecological Sciences, Indian Institute of Science, Bengaluru, India; Département Adaptations du Vivant, UMR 7179, Mécanismes adaptatifs et Évolution (MECADEV) CNRS/Muséum national d’Histoire naturelle, Paris, France; CEFE, Univ Montpellier, CNRS, EPHE, IRD, Montpellier, France; CRNL, Sleep Team, Univ Claude Bernard Lyon1, CNRS, INSERM, Bron, France

**Keywords:** Sleep ecophysiology, Urbanization, Electro-oculogram, EOG, Reptile, Agamidae

## Abstract

Sleep expression in wild animals is regulated by ecological context. Yet, how sleep differs between populations of a species that experience different environmental challenges such as urbanization, remains to be fully understood. Robust quantification of sleep in small animals, such as lizards, has so far been methodologically difficult, limiting population-level comparisons. Using recently developed miniature loggers, we recorded sleep in 19 wild-caught individuals of the South Indian rock agama lizard (*Psammophilus dorsalis*). We compared individuals captured from urban and rural environments and measured electro-oculogram (EOG)-derived sleep parameters: sleep duration, bout frequency, bout duration, and interval between bouts. We then subjected all individuals to an acute stressor (handling-restraint) and examined changes in sleep expression. We found that this species slept mostly at night, with an average total sleep time of 11h 10min. EOG-derived sleep characteristics did not differ substantially between populations, although daytime sleep bouts were more consolidated (shorter interval between bouts) in urban-origin lizards. Contrary to evidence from mammals, acute restraint stress did not impact sleep, overall or for either population. To complement these measures, we also quantified the latency to arousal after exposure to vibration stimuli in a separate set of urban and rural lizards (n = 12 each) and found that urban individuals responded marginally quicker at night compared to rural individuals, demonstrating greater vigilance and/or more fragmented sleep. Overall, we show that sleep characteristics of wild animals in a common garden can be highly conserved within a species, with limited context-dependent variation between populations.

## INTRODUCTION

Sleep occurs across the animal kingdom (Rattenborg and Ungurean, 2023) and is essential for physiological processes, performance, and survival (Aulsebrook et al., 2021; Schmidt, 2014). Sleep expression, however, is variable in terms of duration, distribution, and composition (Rattenborg and Ungurean, 2023), and is responsive to ecological conditions (Capellini et al., 2008; Roth et al., 2006), such as breeding period (Lesku et al., 2012; Zaid et al., 2024), moon phase (van Hasselt et al., 2020), and seasons (Mortlock et al., 2024). Animals also respond to acute stressors, such as predator encounters, by reducing sleep in the aftermath with fewer or shorter sleep bouts (e.g., as observed in wild-caught *Rattus norvegicus*, Lesku et al., 2008; see Tisdale et al., 2018). Such flexibility in sleep expression promotes greater vigilance under risk, enabling animals to respond quicker to threats (Lima et al., 2005). However, current knowledge on the responsiveness of sleep to ecological conditions, including stressors, is limited by taxa and scale, with a dearth of studies on inter-population differences in wild or wild-caught animals and in small vertebrates (Duboué et al., 2011; Libourel et al., 2025).

Urbanization presents a challenging environment for sleeping animals (Lesku and Aulsebrook, 2026), and also provides a test of the context-dependent variation in sleep. In response to urbanization, several behavioural, morphological, and physiological traits shift (Thompson et al., 2022), likely arising from a combination of plasticity, sorting, and contemporary evolution (Bonier, 2023; Ouyang et al., 2019). Of the many environmental axes modified by urbanization, artificial light at night (ALAN) has been shown to result in altered onsets of sleep and awakening, and reduced sleep duration (Aulsebrook et al., 2018; Aulsebrook et al., 2020; Gaston et al., 2017; Raap et al., 2015). Urban environments also have higher noise levels and altered predator assemblages that make sleep more challenging (Connelly et al., 2020; Duarte and Young, 2011; Grunst et al., 2023; but see Caorsi et al., 2019). Moreover, urban life affects stress physiology in some taxa, inducing higher hypothalamic-pituitary-adrenal (HPA) axis activity (e.g., the amplitude of stress response to a stimulus) and altering regulation (e.g., the time taken to revert to baseline levels or ‘decay phase’; Ouyang et al., 2019), which can directly affect sleep (Lo Martire et al., 2020). Overall, animals from urban environments are likely to experience more frequent stimuli while sleeping, and when compounded by the zeitgeber-dilution due to ALAN and potentially altered stress physiology, their sleep duration, consolidation, and distribution could be affected (Dominoni et al., 2013a; Grunst et al., 2023). Although the sleep behaviour of urban and non-urban populations of vertebrates has been compared in some taxa (Mohanty et al., 2021; Ulgezen et al., 2019), we lack robust electrophysiological measures of sleep in individuals across populations.

Sleep expression in response to ecological contexts has been studied with renewed vigour in the last decade, with an integration of behaviour, neurophysiology, and ecology (termed ‘sleep ecophysiology’; Aulsebrook et al., 2016) in several birds and mammals (Aulsebrook et al., 2021; Kendall-Bar et al., 2023; Libourel et al., 2023; Mortlock et al., 2024; Mortlock et al., 2025; Rattenborg et al., 2017). Reptilian sleep and its ecological variations, however, are yet to be examined with such an approach, largely due to methodological constraints of recording small animals (Mohanty et al., 2022). Based on behavioural observations, reptiles have been found to reduce sleep duration under predation risk (Revell and Hayes, 2009; see also Kelly et al., 2015) and respond to urban ALAN conditions with altered sleep site selection (Mohanty et al., 2021) and sleep depth (lowered response latency to stimulus; Kolbe et al., 2021). With the miniaturization of sleep loggers (Massot et al., 2019) and the identification of electro-oculogram (EOG) density as a correlate of lizard sleep (Libourel et al., 2018), sleep ecophysiology can now be extended beyond birds and mammals. This EOG-sleep correlation allows for the quantification of sleep architecture in terms of duration and distribution of sleep bouts (but not sleep “brain state” composition) (Albeck et al., 2022; Blumberg et al., 2020; Libourel et al., 2018).

The South Indian rock agama *Psammophilus dorsalis* is a diurnally active lizard occurring in southern India, with well-known urban ecology (Thaker et al., 2022), stress physiology (Amdekar et al., 2018; Banerjee et al., 2025), active behaviour (Bhave et al., 2017; Radder et al., 2005), and sleep behaviour (Mohanty et al., 2021). Urban individuals show shifts in behaviour, physiology, and even morphology, compared to rural conspecifics (Thaker et al., 2022). Behaviourally, urban lizards are less reactive to human perturbance (lower flight initiation distance; Batabyal et al., 2017) and conspecific interactions (Batabyal and Thaker, 2019) than rural lizards. In terms of physiology, urban lizards exhibit higher baseline and stress-induced circulating corticosterone levels than rural lizards, with a more delayed return to baseline (Batabyal and Thaker, 2019). Yet, broadly, urban individuals fare similarly in terms of most health markers (e.g., body condition, immune response), showing that they largely cope well with urban conditions (Amdekar et al., 2018). Sleep behaviour also shifts in the city, where lizards sleep in sites that partially shelter them from ALAN, likely to avoid direct disruption of sleep by light and/or visibility to predators (Mohanty et al., 2021). Overall, these lizards from urban environments experience higher disturbance conditions that are affecting all aspects of their lives, including sleep. This species, therefore, is ideal to test whether sleep architecture is variable in individuals from urban and rural populations or shows species-level conservation.

Here, we examine population-level differences in sleep expression using wild-caught *P*. *dorsalis* as a model system. We specifically compared urban and rural populations of lizards to determine how prior exposure to greater disturbances experienced by urban lizards and their altered stress physiology would influence sleep expression, sleep response to stress, and arousal response in a common garden condition. First, between the two populations, we compared baseline values of EOG-derived sleep parameters in terms of the percentage of sleep during night and day, the frequency of sleep bouts, as well as the duration of bouts and the interval between bouts. We predicted increased sleep fragmentation (higher frequency, shorter duration, and longer interval between sleep bouts) and greater daytime sleep in urban lizards compared to rural lizards.

The shifted stress physiology of urban animals and the link between stress and sleep may result in urban and rural conspecifics to not only differ in sleep expression but also in the sensitivity of their sleep to stressors. To test this, we subjected the same individuals to an acute stressor to test population differences in sleep’s sensitivity to stress (post-stress vs baseline). Compared to the baseline, we expected lower sleep duration and greater fragmentation of sleep bouts in the first 24-hour period following the stress event, followed by an increase in sleep duration and greater consolidation of sleep bouts in the second 24-hour period post-stress event. For both periods, we expected urban lizards to show a greater magnitude of effect compared to rural lizards, given their shifted stress physiology and regulation. Finally, in a separate set of urban and rural individuals, we compared post-stimulus (vibration) latency to arousal as a behavioural measure of vigilance and sleep depth at night. We expected urban lizards to respond faster than rural lizards to stimuli at night, demonstrating either higher sleep fragmentation and/or increased vigilance. To the best of our knowledge, our study is the first to evaluate interpopulation variation of sleep in wild-caught vertebrates using a robust measure of sleep architecture.

## METHODS

The study was conducted from March to August 2021, coinciding with the activity period of *Psammophilus dorsalis*. We captured only adult male lizards that weighed 75g or more to safely attach ONEIROS-II sleep loggers (Massot et al., 2019) and to not exceed 10% of body weight in attachments. As females of the species weigh less than 50g (Balakrishna et al., 2021), they were excluded from the study. We sampled lizards from three urban sites in Bengaluru city and three rural sites from outside the city (see Mohanty et al., 2021 for details) (**Supporting Information 1 Table 1**). Rural sites had low human disturbance and limited influence of ALAN (lights 200m-1km away from the sampling locations), whereas urban sites were well-illuminated, and were characterised by high human activity, high predation risk (e.g. free-ranging dogs) and vehicular traffic at night. Lizards were captured by lassoing and transported to the lab within four hours of capture.

In captivity, we conducted two experiments on separate sets of individuals. In Experiment 1, we used EOG to compare baseline sleep characteristics between populations, followed by testing the effects of acute stress on sleep, at the species-level and by population origin (10 urban vs 9 rural). In Experiment 2, we compared the latency to arousal during sleep and wake, between urban and rural lizards (12 urban vs 12 rural).

### Care in captivity

In the laboratory, lizards were housed in individual glass terraria (60×30×25 cm) and were exposed to a 12h dark:12h light cycle (dark - 19:00 to 07:00). The terraria were lined with disposable paper towels, covered on all sides to minimize disturbance, and equipped with a rock, a shelter, and petri dishes for food and water. Incandescent basking lights (100 W) above each terrarium were turned on from 1000 to 1100 hours and from 1500 to 1600 hours. Individuals were provisioned daily with mealworms and water, and experienced ambient temperature conditions (24-28°C).

Ethics approval for the study was granted by the Institutional Animal Ethics Committee, Indian Institute of Science (# CAF/Ethics/739/2020).

### Experiment 1 (EOG-derived sleep: effect of population type and stress)

We allowed acclimation to captivity for 3-4 days to reduce the effects of capture and transport stress, while still retaining naturalistic sleep expression. We then attached an ONEIROS-II logger to each lizard to measure EOG and ambient temperature at a sampling rate of 128 Hz (**Supporting Information 1 Fig. 1**). Each lizard was first cold anaesthetized on ice for 5-8 min (Lillywhite et al., 2017) and was placed on an icepack for the procedure. Three stainless steel, fine electrodes (76.2 micron; A-M systems), which record the EOG activity, were placed subcutaneously, two in contact with each eye muscle and one as a reference in the bony portion of the head, just above the neck muscle (adapted for *P. dorsalis* from Libourel et al., 2018). Sites of incision were first numbed with lidocaine hydrochloride and then treated with betadine after electrode positioning. The electrodes were then secured in position by dental acrylic. The logger and battery were positioned onto the back of the lizard with teflon tape and affixed with electrical tape. Lizards were returned to their terraria to recover for 32 hours, after which, the logger was remotely activated and was set to record for two consecutive 24h periods (Baseline Day 1 and Baseline Day 2). On the third day, at 1600h (post-basking), we exposed each individual to an acute stressor by handling it for 15 minutes, during which we changed the battery on the logger, and restraining it in a breathable bag for 45 minutes (Wingfield et al., 1998). Handling-restraint is known to induce a glucocorticoid response in this species and others (Banerjee and Thaker, 2025; French et al., 2006). We then recorded two consecutive 24h periods of post-stress data (Post-stress Day 1 and Post-stress Day 2).

### **Experiment 2** (Arousal response-based vigilance and depth of sleep: effect of population type)

In this experiment, wild-caught lizards (n = 12 urban, 12 rural) were attached with a vibrating micromotor (∼ 5mm in diameter) on the tail base and allowed an acclimation period of 24h in individual terraria. The vibrator was attached by a power cable to an Arduino-based circuit board that allowed us to automate the stimulus intensity and interval between consecutive stimuli. We ensured that the cable was long enough to allow the lizards to move in the terraria without hindrance. The circuit board held a clock and a small LED light that turned on with the vibratory stimulus. Lizards were stimulated by vibrations that lasted for 5 sec every two hours over a 24-hour period. The response of lizards was recorded using two IR-enabled cameras (25 frames/sec, with a precision of 40 msec; Hikvision DS-2CE5ADOT-IRPF). We measured the latency to the first response (eye opening or head and/or body raise or movement) using the video analysis software Tracker (version 5.1.5). We recorded the state of the lizard before the stimulus (‘awake’ or behaviourally ‘asleep’) based on whether the lizard’s eyes were closed and its head and body rested prone on the surface, a typical sleep posture in agamid lizards (Mohanty et al., 2021).

### Data Analysis

#### Experiment 1

EOG information was extracted and processed using custom scripts in MATLAB (R2020b), as per Libourel et al.(2018). Differentials between EOG amplitude between the two eyes were calculated for all individuals (except one, where EOG from only one eye with a reliable signal was used), which was then passed through low-pass (10Hz, order 10) and high pass-filters (2Hz, order 10) and a Hilbert transform. The maximal absolute values were obtained for each second. After visualizing signal quality across individuals, eye movement was defined as a signal with an amplitude of 20 μV or higher (40 μV for the individual with data from one eye). We extracted the number and amplitude of peaks, and the interval between peaks. Then, we scored sleep behaviour similarly to Libourel et al.(2018) and adapted for *P. dorsalis*. Indeed, in the Argentine tegu (*Salvator merianae*), a sharp decrease in eye movements is associated with a high arousal threshold – a marker of sleep, showing that a major shift in eye-movement density can be used to score sleep where immobility alone is insufficient (Libourel et al., 2018). A recent paper shows that in seven reptilian species, eye movement density during behavioural sleep drops substantially compared to wakefulness (Bergel et al., 2025).

We computed the eye movement density from the interval between each eye movement. By looking at the distribution of the intervals between eye movements during behavioural sleep and wake, we considered sleep behaviour when the density was less than 1 eye movement/minute. Two sleep bouts closer than 4 min were merged into one. All other periods were categorised as the awake state, including sleep bouts shorter than 4 min that occurred within wake (**Supporting Information 2)**. Although eye movements may occur during sleep in lizards (Albeck et al., 2022; Bergel et al., 2025), they are often isolated and are likely to subsumed into sleep based on our classification.

We computed, in three-hour and 12-hour groups (coinciding with light-dark phases), four sleep parameters: percentage of sleep, frequency and mean duration of sleep bouts, and the mean interval between sleep bouts. For most analyses, we use data extracted at the 12-hour level (unless stated otherwise) to avoid artificial splitting of sleep bouts. Sleep parameters were extracted for two days each before (Baseline Day 1 and Baseline Day 2) and after acute stress (Post-stress Day 1 and Post-stress Day 2). For the statistical analyses, we only included data from days with complete 24h recordings. Some logger recordings were incomplete due to battery failure and drainage or electrode damage, resulting in varying sample sizes for each objective (**Supporting Information 1 Table 1**). One individual showing high daytime sleep (61.75%) during Baseline Day 1 was discarded from all analyses (**Table 1**).

**Table 1.**
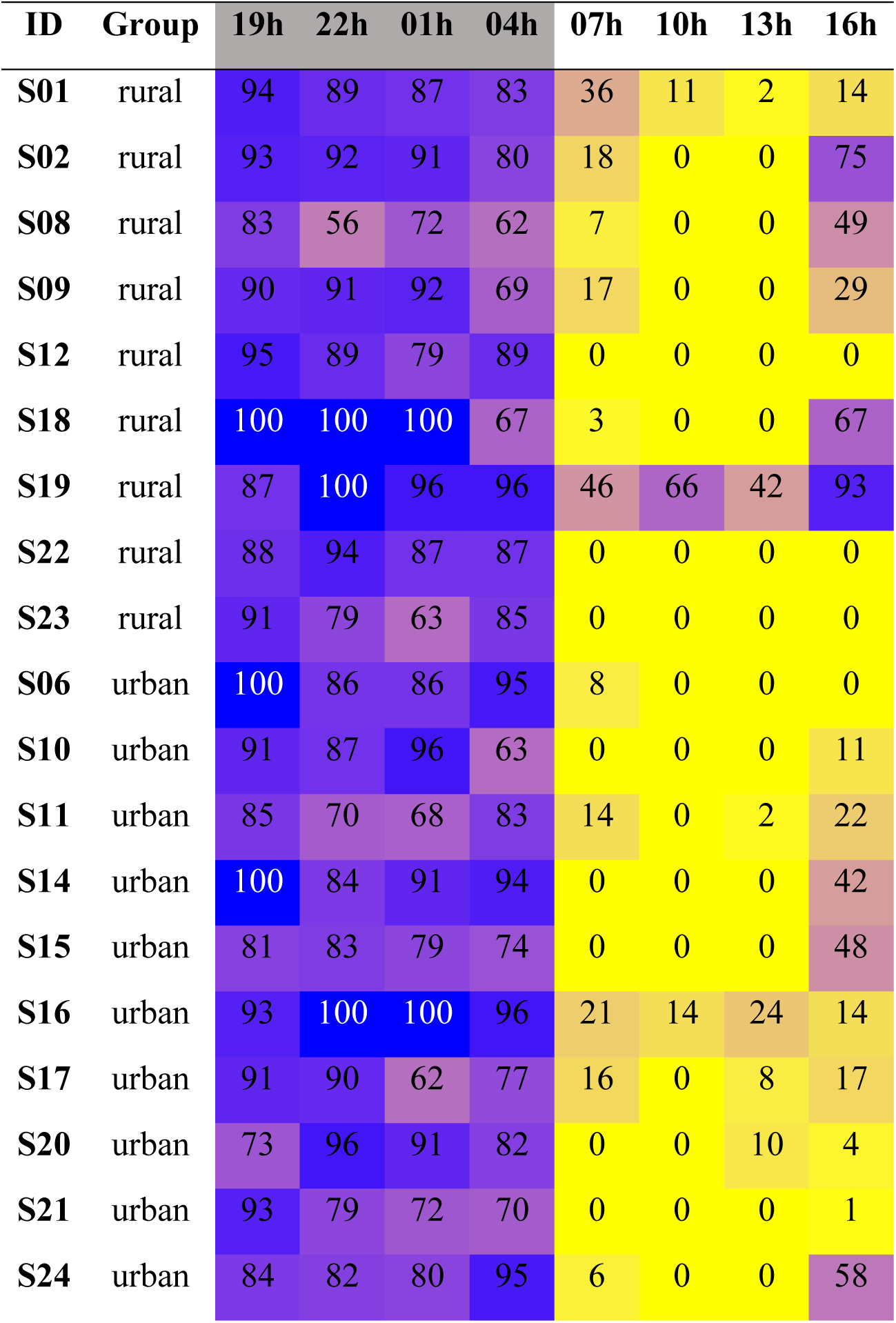
Percentage of sleep in all *Psammophilus dorsalis* individuals tested, shown over three-hour windows starting from 19h (onset of darkness) during Baseline Day 1. Colour gradient shows high values in blue and low values in yellow. Night hours (lights off) are marked in grey. Individual S19 was removed from subsequent statistical analyses.

| ID | Group | 19h | 22h | 01h | 04h | 07h | 10h | 13h | 16h |
| --- | --- | --- | --- | --- | --- | --- | --- | --- | --- |
| S01 | rural | 94 | 89 | 87 | 83 | 36 | 11 | 2 | 14 |
| S02 | rural | 93 | 92 | 91 | 80 | 18 | 0 | 0 | 75 |
| S08 | rural | 83 | 56 | 72 | 62 | 7 | 0 | 0 | 49 |
| S09 | rural | 90 | 91 | 92 | 69 | 17 | 0 | 0 | 29 |
| S12 | rural | 95 | 89 | 79 | 89 | 0 | 0 | 0 | 0 |
| S18 | rural | 100 | 100 | 100 | 67 | 3 | 0 | 0 | 67 |
| S19 | rural | 87 | 100 | 96 | 96 | 46 | 66 | 42 | 93 |
| S22 | rural | 88 | 94 | 87 | 87 | 0 | 0 | 0 | 0 |
| S23 | rural | 91 | 79 | 63 | 85 | 0 | 0 | 0 | 0 |
| S06 | urban | 100 | 86 | 86 | 95 | 8 | 0 | 0 | 0 |
| S10 | urban | 91 | 87 | 96 | 63 | 0 | 0 | 0 | 11 |
| S11 | urban | 85 | 70 | 68 | 83 | 14 | 0 | 2 | 22 |
| S14 | urban | 100 | 84 | 91 | 94 | 0 | 0 | 0 | 42 |
| S15 | urban | 81 | 83 | 79 | 74 | 0 | 0 | 0 | 48 |
| S16 | urban | 93 | 100 | 100 | 96 | 21 | 14 | 24 | 14 |
| S17 | urban | 91 | 90 | 62 | 77 | 16 | 0 | 8 | 17 |
| S20 | urban | 73 | 96 | 91 | 82 | 0 | 0 | 10 | 4 |
| S21 | urban | 93 | 79 | 72 | 70 | 0 | 0 | 0 | 1 |
| S24 | urban | 84 | 82 | 80 | 95 | 6 | 0 | 0 | 58 |

For each of the four EOG-derived sleep parameters in Experiment 1, we built two linear mixed-effects models. The approach of two separate models was chosen to be consistent with our hypotheses, where the first model included data from the two Baseline days (Baseline Day 1 & 2) to investigate the differences in baseline sleep between populations, whereas the second model encompassed all four days (Baseline Day 1 & 2 and Post-stress Day 1 & 2) to test stress-induced changes to sleep. The models with only Baseline days allowed us to assess the stability of sleep parameters across the two Baseline days. Following this, we used the mean of both baseline days coded by a planned contrast in the other model (see **Supporting Information 1**). The second model with 2 Baseline and 2 Post-stress days tested changes in sleep characteristics following the stressor, by comparing the average of Baseline days to Post-stress day 1 and Post-stress day 2. Both sets of models included lizard ID as a random effect, and as fixed effects the day of experiment (Baseline Day 1 & 2, Post-stress Day 1 & 2), population origin (‘urban’, ‘rural’), and diel phase (‘daytime’, ‘nighttime’). We also included the mean ambient temperature of each phase as a covariate. For both sets of models, we modelled interaction terms between diel phase and population type, as well as the day of experiment and population type.

Of the four sleep parameters as response variables, we used a negative binomial generalized linear mixed model for frequency of sleep bouts (using the ‘glmmTMB’ package); the other three were modelled as Gaussian variables (using the ‘lmerTest’ package). Two response variables, duration and interval between sleep bouts, were log-transformed. We coded categorical predictors as sum-to-zero contrasts prior to model building (Fox and Weisberg, 2019) and used type III tests, i.e., type III Wald χ² for frequency of sleep bouts and type III Anova tests (with Kenward-Roger’s method) for the other three variables, using the package ‘car’. For all models, we first evaluated statistical significance of interaction terms and only significant interactions were subjected to post-hoc comparisons using the ‘emmeans’ package with Holm p-value adjustment. Independently, our planned contrasts compared the average of Baseline days vs Post-stress day 1, and the average of Baseline days vs Post-stress day 2. Additionally, we evaluated the immediate effect of acute stress on *P. dorsalis* sleep by using EOG-derived parameters calculated at the three-hour level. These models included lizard ID as a random effect, and as fixed effects the day of experiment (Baseline Day 1 & 2, Post-stress Day 1) and hour-group (e.g., ‘19h-22h’, ‘22h-01h’ etc), including an interaction term. We also included mean ambient temperature of each hour-group as a covariate.

#### Experiment 2

For the data from experiment 2, we first considered observations only when individuals were behaviourally asleep at night (defined above), leading to one individual being removed (n = 12 rural, 11 urban); no individuals were behaviourally asleep during daytime. All individuals responded to the vibration stimulus with arousal (opening eyes or moving limbs and neck). As individuals received 6 stimuli at night, each separated by 2 hours, we also checked if the number of stimuli previously received affected the arousal response. We built a GLMM with log-transformed latency to arousal as the response, stimulus number (1^st^-6^th^) and population origin (‘urban’, ‘rural’) as fixed effects, and lizard ID as a random effect. To investigate response latency according to diel phase, irrespective of behavioural sleep, we built a GLMM with diel phase (‘daytime’, ‘nighttime’) interacting with population origin, along with stimulus number over both daytime and nighttime (1^st^-12^th^) as fixed effects, and lizard ID as a random effect. Significant interactions were further tested using post-hoc comparisons in the ‘emmeans’ package with Holm p-value adjustment.

Data exploration was carried out following Zuur et al. [56]. We checked for outliers in the data and built generalized mixed models; model fitting and reporting followed Zuur & Ieno (2016). All models were evaluated for validity including model residual distribution, overdispersion, and zero-inflation (for frequency of sleep bouts), using the ‘DHARMa’ package (Hartig, 2026). We quantified model fit based on R^2^_GLMM_ (Nakagawa et al., 2017). All analyses were carried out in R (R Core Team 2026).

## RESULTS

### Baseline sleep in Psammophilus dorsalis (Experiment 1a)

At the species-level on Baseline days, *P. dorsalis* slept for an average of 11h 10 min (SE = 21.3min; range: 7h 23min – 13h 55min; n = 18). Sleep occurred predominantly at nighttime (**Table 1**) and on average, spanned 84.1% of the night (SE = 2.04; range: 61.5 – 97.7), distributed over 13.3 bouts (SE = 1.35; range: 3 – 24) of 1h 8min each (SE = 15.18min; range: 19.41min – 4h 5min) which were separated by an interval of 8.11 min (SE = 0.35min; range: 6.23 – 10.93min; **Table 2**). In contrast, during the day, lizards on average slept only 8.89% of the time, over 4.53 short bouts of 17.23min, with a gap of 1h 52min between them (**Table 2**). Sleep parameters did not differ between the two Baseline days (**Supporting Information 1 Table 2**).

**Table 2.**
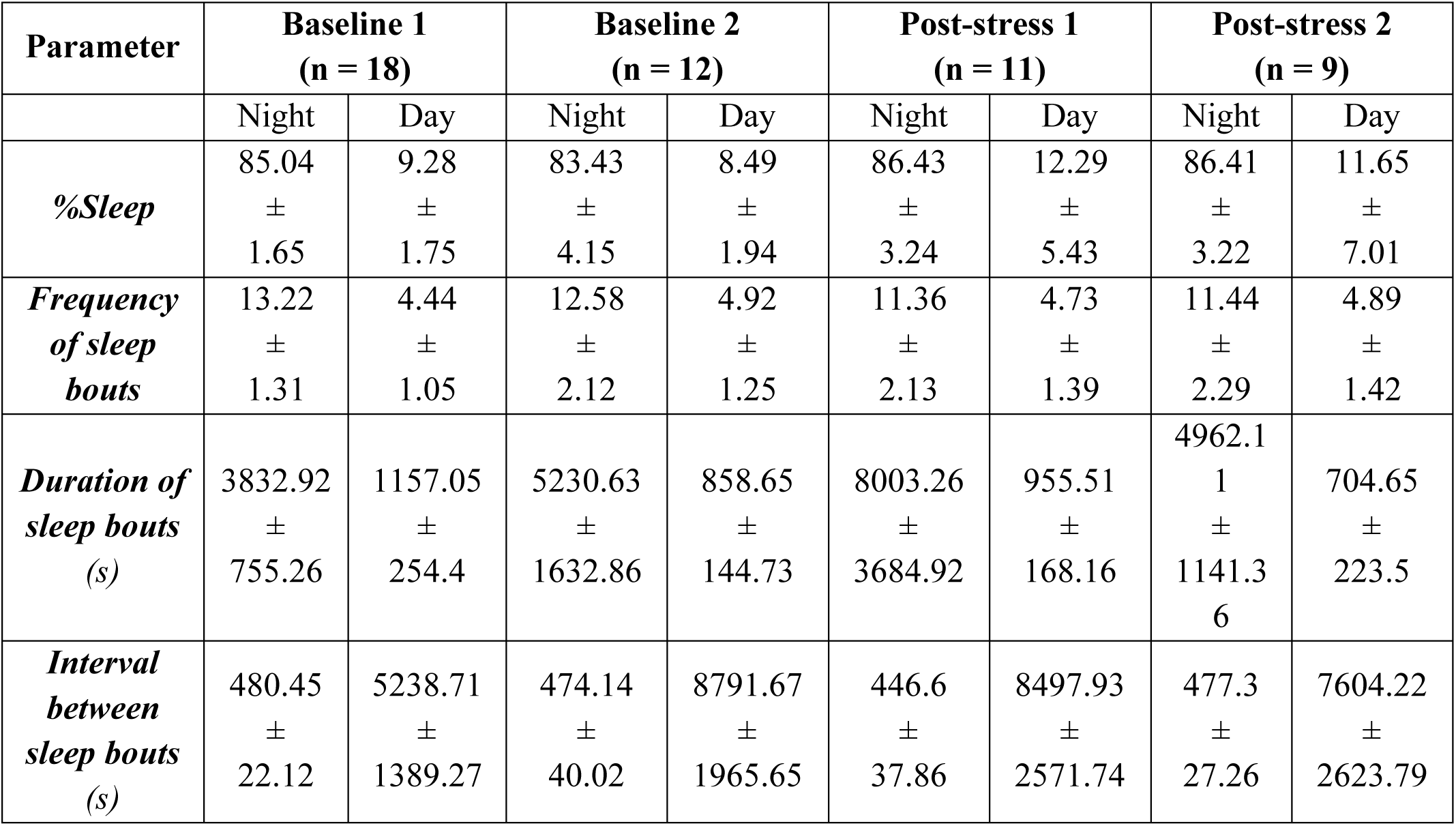
EOG-derived sleep parameters in *Psammophilus dorsalis* (n = 18) across four 24h periods: baseline (Day 1 and Day 2) and post-stress (Day 1 and Day 2). Except %Sleep, all parameters pertain to sleep bouts. Values are mean ± SE.

Overall, rural and urban lizards did not differ in any sleep parameter on Baseline days (**Supporting Information 1 Table 2, Fig. 2**), after accounting for diel phase and ambient temperature. The interaction between diel phase and population origin was also not significant for any sleep parameter (**Supporting Information 1 Table 2**) but was marginally significant in the case of the interval between sleep bouts (*F =* 3.94; *p = 0.054*). A contrast using estimated marginal means showed that during daytime, mean intervals between sleep episodes were approximately 2.2 times longer in rural than urban lizards (Estimate ± SE = 0.787 ± 0.335 on the log scale, *p* = 0.025).

### Inter-population variation in sleep response to acute stress (Experiment 1b)

#### 24h and 48h after an acute stress event

After accounting for diel phase and ambient temperature, sleep parameters did not differ on Post-stress Day 1 & 2 in relation to the Baseline days (**Fig 2**, **3**; **Table 3**). Planned contrasts comparing Post-stress Day 1 & 2 to the average of the Baseline days were likewise non-significant (**Supporting Information 1 Table 4**). Population origin had no significant interaction effect with the day of experiment for all sleep parameters (**Table 3**). However, over all four days of the experiment, urban lizards had approximately 2.4-fold shorter mean intervals between sleep bouts than rural lizards during daytime.

**Figure 1.**
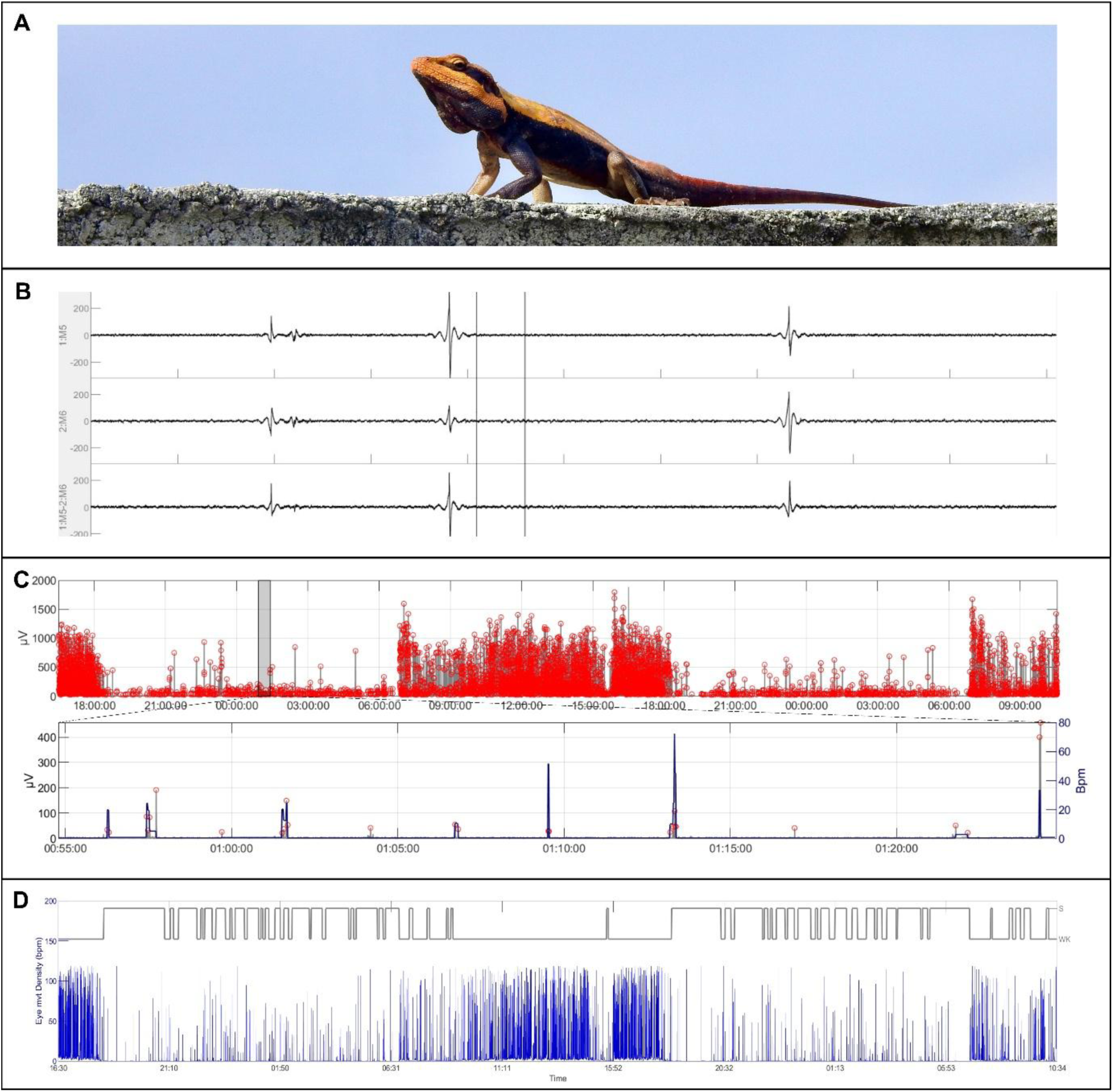
Sleep quantification using electro-oculograms (EOG) in (A) the Peninsular rock agama *Psammophilus dorsalis* (Photo: Nimish Anil). (B) Raw EOG signals of left eye, right eye, and the differential (from top to bottom), (C) Algorithmic detection of eye movement (with a zoomed interval), (D) Classification of eye movement density into bouts of sleep (S) and wake (WK).

**Figure 2.**
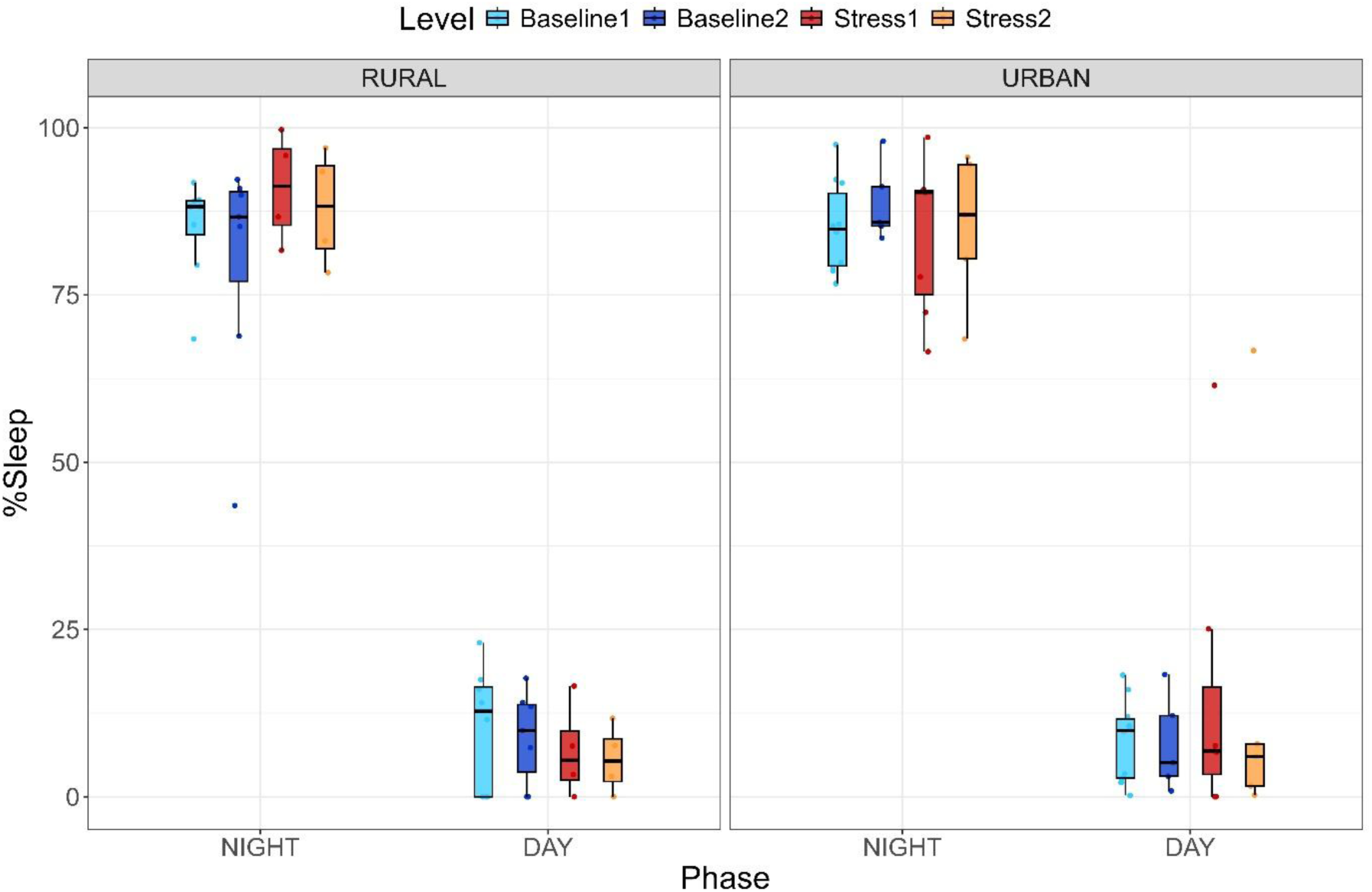
Variation in EOG-derived percentage of sleep of rural and urban lizards in response to acute stress event on Post-stress Day 1 (red) and Post-stress Day 2 (orange), with respect to Baseline Day 1 (light blue) & Baseline Day 2 (dark blue).

**Figure 3.**
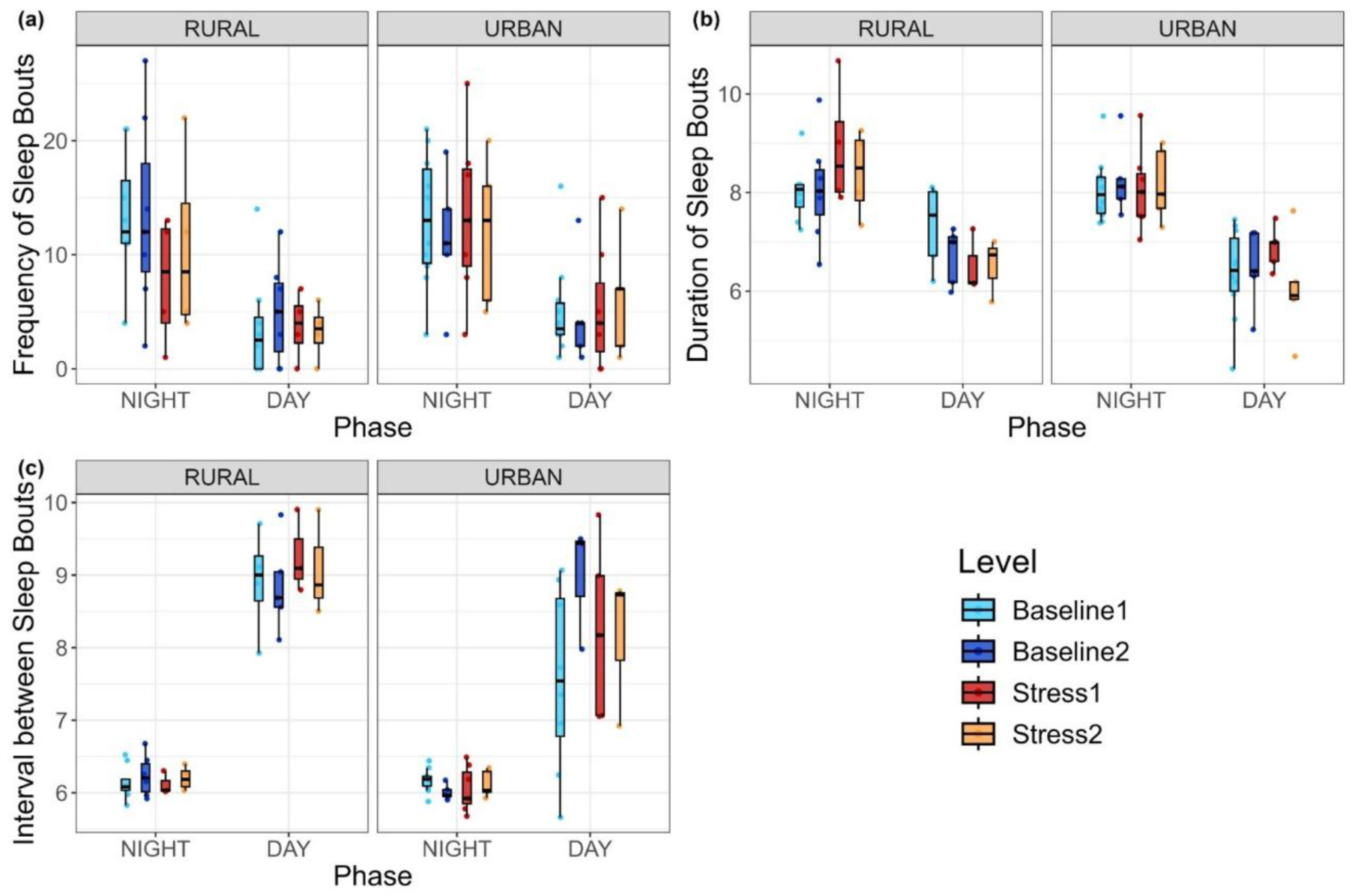
Response of EOG-derived sleep bout parameters (*a*-*c*) of rural and urban lizards to acute stress event on Post-stress Day 1 (red) and Post-stress Day 2 (orange), with respect to Baseline Day 1 (light blue) & Baseline Day 2 (dark blue).

**Table 3.** The influence of diel phase (night and day) and population origin (urban and rural) on four sleep parameters, on all four days (Baseline Day 1 and 2, Post-stress Day 1 and 2). All models include the model components as fixed effects, along with lizard ID as a random effect. Statistics reported are from linear mixed models with Type III Wald χ² or Anova tests (with Kenward-Roger’s method), using sum-to-zero contrasts.

| **Model component** | ***F (or $\chi^2$)*** | ***p*** | ***R<sup>2</sup><sub>m</sub>*** | ***R<sup>2</sup><sub>c</sub>*** |
| --- | --- | --- | --- | --- |
| ***%Sleep*** |  |  | 0.90 | 0.94 |
| Day | 0.30 | 0.82 |  |  |
| Phase | 1217.65 | **<0.001** |  |  |
| Origin | 0.55 | 0.469 |  |  |
| Day x Origin | 0.77 | 0.509 |  |  |
| Phase x Origin | 0.49 | 0.485 |  |  |
| Temperature | 0.20 | 0.652 |  |  |
| ***Frequency of sleep bouts*** |  |  | 0.37 | 0.39 |
| Day | 0.52 | 0.913 |  |  |
| Phase | 36.66 | **<0.001** |  |  |
| Origin | 1.51 | 0.218 |  |  |
| Day x Origin | 0.67 | 0.878 |  |  |
| Phase x Origin | 0.77 | 0.377 |  |  |
| Temperature | 1.96 | 0.161 |  |  |
| ***Duration of sleep bouts*** |  |  | 0.51 | 0.65 |
| Day | 0.47 | 0.699 |  |  |
| Phase | 92.96 | **<0.001** |  |  |
| Origin | 0.27 | 0.605 |  |  |
| Day x Origin | 0.25 | 0.861 |  |  |
| Phase x Origin | 0.33 | 0.562 |  |  |
| Temperature | 0.07 | 0.781 |  |  |
| ***Interval between sleep bouts*** |  |  | 0.76 | 0.79 |
| Day | 0.74 | 0.528 |  |  |
| Phase | 240.84 | **<0.001** |  |  |
| Origin | 4.96 | **0.044** |  |  |
| Day x Origin | 0.70 | 0.553 |  |  |
| Phase x Origin | 8.15 | **0.005** |  |  |
| Temperature | 0.24 | 0.629 |  |  |

We also tested the immediate response of lizards after the stress event (at 3h-level instead of 12h). Across all four EOG-derived sleep parameters, there was no evidence of an acute effect of the stress event during the first 3h of the dark phase, and no evidence that the stress effect varied across the subsequent 3h intervals (**Supporting Information 1 Table 5**).

### Inter-population variation in sleep depth/latency to arousal response (Experiment 2)

At night, behaviourally asleep urban lizards responded quicker to stimuli (β = -0.36, SE = 0.16, *p =* 0.038; **Fig. 4**) than rural lizards. We found no significant effect of stimulus number (*β* = 0.01; *p =* 0.599). However, model performance was weak (R^2^_m_ = 0.09) with a strong random effect (R^2^_c_ = 0.37). Confidence intervals overlapped between rural (247 – 399msec) and urban lizards (172 – 278msec).

**Figure 4.**
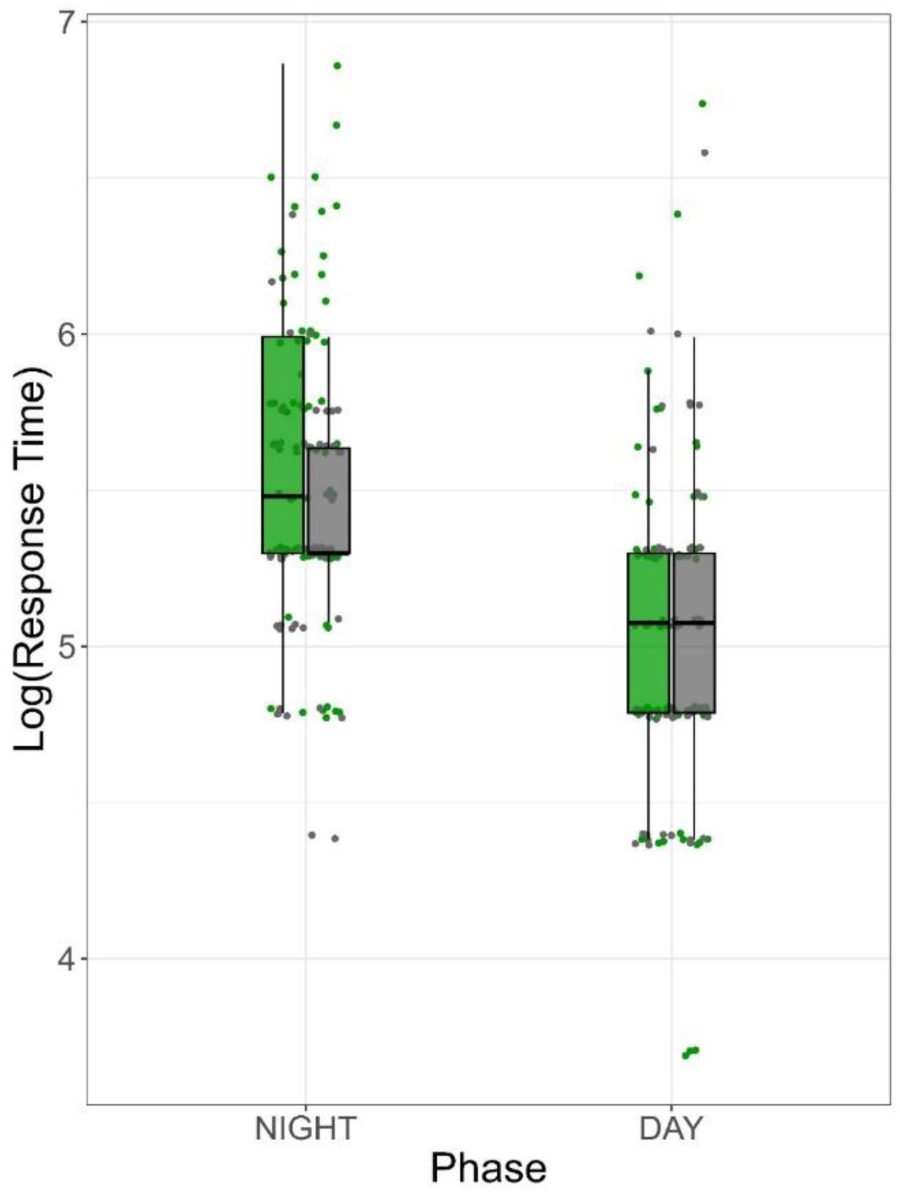
Latency to arousal after vibration stimulus shown by rural (green) and urban (grey) lizards during the day and night.

We also compared the response latency to a stimulus between daytime and nighttime, across populations. We found a significant interaction between diel phase and population (β = 0.23, SE = 0.11, *p =* 0.039, R^2^_m_ = 0.19, R^2^_c_ = 0.24). All lizards showed quicker responses during daytime than nighttime (β = -0.60, SE = 0.09, *p <* 0.001; **Fig. 4**), but urban lizards showed quicker responses than rural lizards only at night (*t* = 2.20; *p =* 0.031).

## DISCUSSION

The variable expression of sleep in a species under different ecological contexts, is not only of fundamental interest but can also influence how species respond to global change phenomena such as urbanization. Although sleep phenotypes have been compared between congeneric species (Joyce et al., 2024), little is known about inter-population differences, such as those between urban and rural conspecifics (in humans; Smit et al., 2019; Yetish et al., 2015). This study provides the first such comparison of EOG-derived sleep parameters in wild-caught vertebrates, sampled from multiple sites within urban and rural areas. Contrary to expectations, we found substantial conservation of sleep phenotype in *Psammophilus dorsalis* lizards, in common garden conditions. Urban-origin individuals showed only minor differences in their sleep by consolidating daytime sleep bouts more than rural-origin individuals, and by slightly decreasing their latency to arousal when stimulated. Sleep in these lizards also displayed high resilience to acute stress in contrast to what has been reported for rodents and humans (Lo Martire et al., 2020). Similar to other species, we found that sleep phenotypes vary considerably between individuals.

Overall, *P*. *dorsalis* lizards slept mostly at night, across 13 sleep bouts on average. However, daytime sleep was not negligible (64 ± 12 min), being concentrated in the hours leading up to darkness (16h-19h; **Table 1**). This consolidated sleeping pattern in accordance with the light phase has been observed in other diurnally-active lizard species, such as the Common green iguana (*Iguana iguana*; Ayala-Guerrero and Mexicano 2008), the Egyptian rock agama (*Laudakia vulgaris*; Albeck et al. 2022), the Bearded dragon (*Pogona vitticeps*; Shein-Idelson et al. 2016) and the Argentine tegu (*Salvator merianae*; Libourel et al., 2018). Lizard sleep is likely governed mechanistically by circadian rhythms, and maintained by the natural light cycle (Yamaguchi et al. 2024). Compared to the day, sleep in *P. dorsalis* was distributed in relatively long and numerous bouts with shorter intervals between them at night. The short wake intervals between sleep bouts at night could serve a vigilance role, as active and passive environmental awareness during the sleep phase is an important anti-predator strategy in reptiles (Lima et al., 2005; Mohanty et al., 2022). Finally, we found high inter-individual variation in sleep parameters, such as the duration of sleep bouts at night, suggesting that despite similar lab conditions, individuals vary in sleep characteristics.

Sleep expression in the lab was generally consistent across urban and rural populations, with no changes in sleep duration and frequency of sleep bouts. Our findings contrast with the only other quantitative inter-population comparison of sleep-like behaviour, observed between populations of the Characin fish *Astyanas mexicanus* (Duboué et al., 2011), where cave populations expressed 14-31% of the sleep duration of surface populations. We observed consolidated bouts during daytime sleep in urban lizards, which may allow more time for vigilance during the rest of the day (Lima et al., 2005), especially as urban individuals encounter a greater frequency of stimuli in their environment. Similarly, urban lizards responded marginally quicker to vibration stimulus at night; this suggests that the depth of sleep is shallower in urban lizards compared to rural lizards, which could bolster vigilance. Quicker arousal to stimuli at night could also be an indicator of poor sleep. In response to short-term exposure to ALAN in the wild, the Cuban brown anole *Anolis sagrei* lizards show quicker arousal from behaviourally-defined sleep (Kolbe et al. 2021). Yet in our study, wild-caught, urban *P. dorsalis* lizards with long-term exposure to ALAN did not vary substantially in arousal response from rural populations (only 9% of variation in arousal latency was accounted for by the population effect) when brought into a common garden condition. This indicates that urbanization-associated ALAN effect on arousal response is either limited to shorter timescales or is plastically expressed. Apart from increased encounters of stimuli in urban areas, a weak zeitgeber effect due to ALAN may lead to changes in urban sleep architecture and onset (i.e., sleep less consolidated by diel phase; Dominoni et al., 2013b). Yet, we did not observe such an increase in sleep during the day by urban lizards. Overall, the observed lack of strong population-level differences in the sleep phenotype of these lizards suggests conservation of sleep characteristics despite differences in prior ecological contexts. A lack of variability in sleep might be maladaptive to the requirements of living in urban environments.

Contrary to expectations, we did not observe an effect of acute stress on sleep architecture. Lizards showed no change in sleep parameters, either in the short-term (between 2 to 5 hours following the stressor) or long-term (in the two 24h periods post-stress; Figs. 2, 3). Our findings are contrary to the suppressed sleep (and increased vigilance) observed in wild-caught rodents the first 3 hours after a simulated predator attack (Lesku et al. 2008). In captive pigeons, an increase in perceived risk alters sleep state composition but does not change the overall sleep duration (Tisdale et al. 2018). Although predator presence has been shown to reduce behaviourally-measured sleep in lizards (*Dipsosaurus dorsalis;* Revell and Hayes 2009), we do not find this pattern in *P*. *dorsalis*. It is possible that the handling and restraining protocol we employed is not equivalent to direct predator encounters, as type, duration, intensity, predictability, and controllability of stressors can affect sleep in different ways (Sanford, Suchecki, and Meerlo 2015; Lo Martire et al. 2020). Similarly, Morlock et al. (2025) found no impact of early-life, chronic stress, measured from hair cortisol, on sleep-like behaviour of neonate fallow deer *Dama dama* in the wild. The generality of stress-sensitive sleep, in response to acute or chronic stress, needs to be evaluated more broadly.

We find a similar lack of stress-induced change in sleep architecture in urban and rural lizards, despite known differences in stress physiology between these populations (i.e., higher baseline corticosterone and longer decay time in urban lizards; Batabyal and Thaker, 2019). Following 30 min of social interactions, corticosterone concentrations in males of *P. dorsalis* show a peak at 20 min post-interaction, followed by a steady reduction in levels from 30-120min post-interaction (Batabyal and Thaker 2019). Even with a longer period of stress induction in this experiment (ca. 60 min; 16h-17h), it is possible that stress response peaked much before normal sleep onset (19h, lights off) and hence, did not ultimately hamper sleep expression. Alternately, the absence of change in sleep characteristics after acute restraint stress might also support the conclusion that sleep is fairly conserved rather than highly flexible. Overall, for urban populations of *P. dorsalis*, behavioural and physiological responses (including sleep) to urban conditions may be decoupled (Koolhaas et al. 2011; Guindre-Parker, Kilgour, and Linkous 2022). For example, the observed plasticity in sleep site choice in urban and rural lizards (Mohanty et al., 2021) may allow for expression of similar sleep architecture. Replicating sleep measurements in the wild is necessary to assess whether sleep architecture differs from the patterns observed in captivity. Furthermore, approaches in experimental evolution that simultaneously measure multiple behavioural and physiological responses can elucidate the genetic and environmental mechanisms underlying the urban phenotypes of these lizards (Ouyang et al., 2019).

Although our experiments tested the effects of urbanization and acute stress with robust sleep measurements, the inferences we draw must be tempered with some considerations. Studying sleep in captivity, while providing a common garden condition for different populations, can lead to patterns not representative of wild sleep due to captivity-induced stress (even in a short period of time) and low enrichment and perceived risk (Rattenborg et al. 2017). The attachment of the logger itself could have impacted the baseline data (especially Baseline Day 1) even 32 hours later, which might contribute to the observed similarity in sleep before and after an acute-stress event. Although EOG allowed us to quantify sleep in several individuals within a short period of time, some methodological aspects require further optimization before it can be widely used (see **Supporting Information 1**). Captivity and surgery effects can also be remedied in future experiments with mesocosm conditions and greater acclimation time.

Another key aspect is the effect of sex on sleep that we could not test due to technological constraints, as females were too small for these loggers. *Psammophilus dorsalis* sexes diverge in morphology, habitat use, behaviour, and evolution of traits (Balakrishna et al., 2021; Parambil and Isvaran, 2025; Ranade and Isvaran, 2022), and so it is possible that unlike males, sleep in females could be more sensitive to environmental contexts and stress.

Research on sleep ecophysiology has made rapid progress in the last decade by quantifying sleep phenotypes in several hitherto unrecorded taxa and testing ecological context-dependence (Kendall-Bar et al., 2023; Libourel et al., 2023; Loftus et al., 2024; Mortlock et al., 2024; Mortlock et al., 2025; Rattenborg et al., 2016; Zaid et al., 2024). These studies indicate flexibility of sleep expression in response to short-term ecological needs and variability (to an extent) between individuals and populations. Although in this study, under captive conditions, wild-caught lizards from urban and rural populations exhibited similar sleep phenotypes, considering inter-individual sleep variability could elucidate the role of sleep in an animal’s response to environmental challenges. Whether the inability of some species to express context-specific sleep phenotypes makes them vulnerable to global change needs to be evaluated. Our results also do not support any effects of acute stress on sleep, showing the need to consider a wide range of vertebrate taxa, including reptiles, in sleep research. Direct tests of ecological context-dependence (e.g., under high predation risk, anthropogenic disturbance) of sleep in the wild or in mesocosm experiments with large sample sizes would further our understanding of the flexibility and variability of sleep.

## Acknowledgments

We thank members of the Macrophysiology Lab for help with animal sampling and captive care, and Anthony Herrel for enabling collaborations between the authors. We are grateful to Siddharth Nair and Kavya for building the arousal response setup. This research was funded by the Indian Institute of Science’s Raman Postdoctoral Fellowship (to NPM), European Union’s Horizon research and innovation programme under the Marie Skłodowska-Curie grant (#101108046 to NPM), and the DBT-Wellcome Trust India Alliance Grant (IA/I/19/2/504639; to MT).

