## Supporting Information 1 for "How variable and stress-sensitive is sleep expression in lizards?"

**SI Fig. 1** Miniature logger implantation on *Psammophilus dorsalis* lizard.

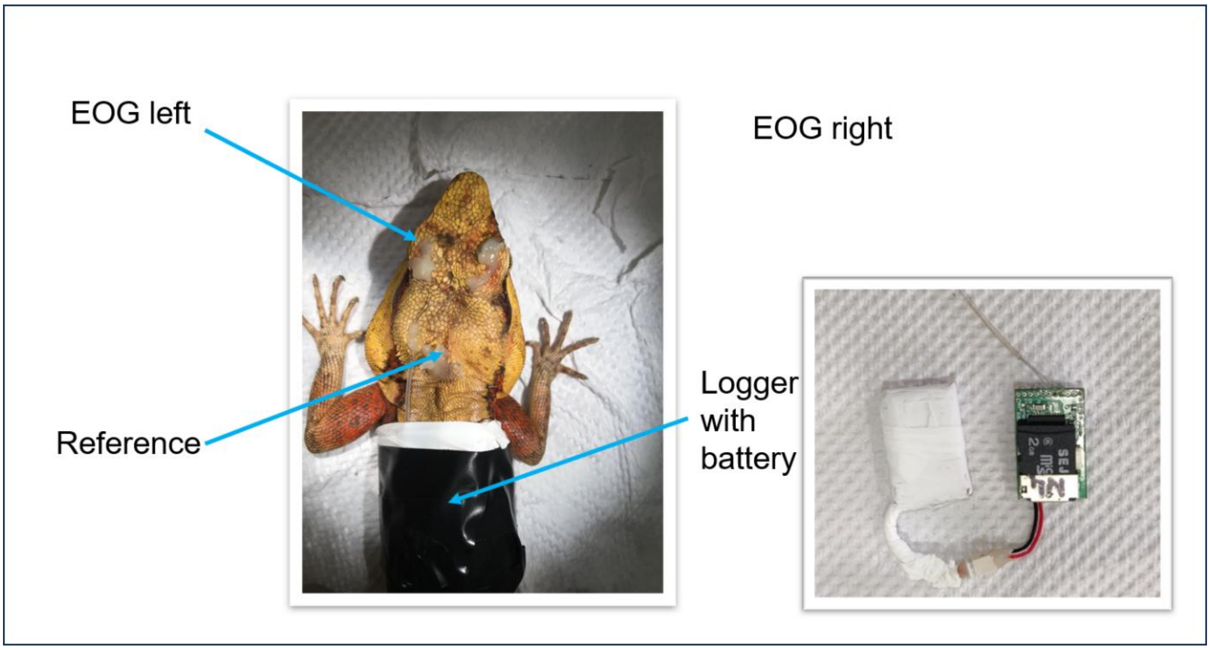

**Limitations of EOG-based sleep scoring**: The use of EOG as a correlate of sleep, though demonstrated in the case of *Salvator merianae*, requires validation in *P. dorsalis* by comparing EOG, arousal response, and sleep posture in the same individuals. Subtle differences among populations in neurophysiology and varying vigilance associated with brain states cannot be detected. Eye movements may also occur during sleep which could influence the classification of sleep bouts. The use of threshold-based sleep scoring, though useful for intra-specific studies, can hinder direct comparisons of bout duration and frequency between species and therefore must be carefully considered.

**SI Table 1.** Sample sizes of *Psammophilus dorsalis* lizards from six urban sites and rural sites in and around Bengaluru city.

| Population type | Site | Experiment 1 | Experiment 2 |
|---|---|---|---|
| Rural | Avati | 3 | 5 |
| | Kolar 1 | 3 | 4 |
| | Kolar 2 | 3 | 3 |
| Urban | Sahakarnagar 1 | 4 | 3 |
| | Sahakarnagar 2 | 4 | 4 |
| | Ullal | 2 | 5 |

**SI Table 2.** Data completeness by individuals across four days of the experiment (Baseline Day 1 & 2; Post-stress Day 1 & 2).

| ID | Origin | Baseline1 | Baseline2 | Stress1 | Stress2 |
|---|---|---|---|---|---|
| **S01** | Rural | ✓ | ✓ | - | - |
| **S02** | Rural | ✓ | ✓ | ✓ | ✓ |
| **S08** | Rural | ✓ | ✓ | - | - |
| **S09** | Rural | ✓ | ✓ | ✓ | ✓ |
| **S12** | Rural | ✓ | ✓ | ✓ | ✓ |
| **S18** | Rural | ✓ | ✓ | ✓ | ✓ |
| **S19** | Rural | ✓ | ✓ | ✓ | ✓ |
| **S22** | Rural | ✓ | - | - | - |
| **S23** | Rural | ✓ | ✓ | - | - |
| **S06** | Urban | ✓ | ✓ | - | - |
| **S10** | Urban | ✓ | ✓ | ✓ | ✓ |
| **S11** | Urban | ✓ | - | ✓ | - |
| **S14** | Urban | ✓ | ✓ | ✓ | ✓ |
| **S15** | Urban | ✓ | - | ✓ | - |
| **S16** | Urban | ✓ | ✓ | ✓ | ✓ |
| **S17** | Urban | ✓ | - | ✓ | ✓ |
| **S20** | Urban | ✓ | - | - | - |
| **S21** | Urban | ✓ | ✓ | ✓ | ✓ |
| **S24** | Urban | ✓ | - | - | - |
| **Total** | | **19** | **13** | **12** | **10** |

**SI Table 3.** The influence of diel phase (night and day) and population origin (urban and rural) on four sleep parameters, from on **baseline** days (Baseline Day 1 and 2). All models include the model components as fixed effects, along with lizard ID as a random effect. Statistics reported are from linear mixed-effect models with Type III Wald $\chi^2$ or Anova tests (with Kenward-Roger's method), using sum-to-zero contrasts.

| **Model component** | ***F (or $\chi^2$)*** | ***p*** | $R^2_m$ | $R^2_c$ |
|---|---|---|---|---|
| ***%Sleep*** | | | 0.94 | 0.97 |
| Day | 0.49 | 0.483 | | |
| Phase | 1465.59 | **<0.001** | | |
| Origin | 0.16 | 0.692 | | |
| Day x Origin | 1.33 | 0.254 | | |
| Phase x Origin | 2.26 | 0.140 | | |
| Temperature | 0.02 | 0.872 | | |
| ***Frequency of sleep bouts*** | | | 0.41 | 0.41 |
| Day | 0.07 | 0.788 | | |
| Phase | 26.07 | **<0.001** | | |
| Origin | 0.23 | 0.629 | | |
| Day x Origin | 0.17 | 0.679 | | |
| Phase x Origin | 0.72 | 0.395 | | |
| Temperature | 2.50 | 0.113 | | |
| ***Duration of sleep bouts*** | | | 0.49 | 0.60 |
| Day | 0.009 | 0.921 | | |
| Phase | 51.37 | **<0.001** | | |
| Origin | 0.65 | 0.430 | | |
| Day x Origin | 0.85 | 0.360 | | |
| Phase x Origin | 3.03 | 0.089 | | |
| Temperature | 0.77 | 0.390 | | |
| ***Interval between sleep bouts*** | | | 0.73 | 0.73 |
| Day | 1.41 | 0.241 | | |
| Phase | 114.33 | **<0.001** | | |
| Origin | 2.88 | 0.111 | | |
| Day x Origin | 1.13 | 0.294 | | |
| Phase x Origin | 3.94 | 0.054* | | |
| Temperature | 0.22 | 0.641 | | |

**Additional modelling rationale**:

The Baseline days only models allowed testing the stability of sleep parameters on the two Baseline days, without necessitating such comparisons across all days, i.e., without comparing Post-stress Day 1 to Baseline Day 1, PS1 to B2, PS2 to B1, and PS2 to B2, which leads to multiple unutilised contrasts. We did not include an interaction term between the day of experiment and diel phase as we did not have an *a priori* hypothesis; similarly, we excluded a three-way interaction to reduce parameter burden. In doing so, we tested the stated hypotheses with a statistically disciplined approach, i.e., no unnecessary interactions or comparisons that increase parameter burden, for our moderate sample size (n = 18 individuals).

**SI Table 4.** Planned contrasts from linear mixed models comprising all four days of the experiment, using sum-to-zero contrasts. Sleep parameters are compared between Post-stress Day 1 (denoted as Stress1) or Post-stress Day 2 (Stress2) and average of the Baseline days (Baseline).

| Sleep parameter | Contrast | Estimate | SE | *t/z* | Adjusted *P* |
|---|---|---|---|---|---|
| ***%Sleep*** | Baseline vs Stress1 | -1.53 | 2.44 | -0.62 | 1.000 |
| | Baseline vs Stress2 | -0.87 | 2.57 | -0.34 | 1.000 |
| ***Frequency of sleep bouts*** | Baseline vs Stress1 | 0.13 | 0.18 | 0.69 | 0.969 |
| | Baseline vs Stress2 | 0.04 | 0.19 | 0.23 | 0.969 |
| ***Duration of sleep bouts*** | Baseline vs Stress1 | -0.14 | 0.20 | -0.69 | 0.950 |
| | Baseline vs Stress2 | 0.14 | 0.20 | 0.71 | 0.950 |
| ***Interval between sleep bouts*** | Baseline vs Stress1 | -0.06 | 0.18 | -0.35 | 1.000 |
| | Baseline vs Stress2 | -0.03 | 0.19 | -0.17 | 1.000 |

**SI Table 5.** Mixed-effects models examining the effect of an acute stress event on EOG-derived sleep parameters during the first 3 h of the dark phase (19:00–22:00; lights off at 19:00; stress event at 16:00–17:00), comparing Post-stress Day 1 with Baseline Day 1 during the corresponding period. Sleep parameters during subsequent 3-h intervals on Post-stress Day 1 also did not differ from the corresponding intervals on Baseline Day 1 (not shown). *Singular fit; the estimated variance of the individual-identity random effect was zero.

| **Parameter** | **Estimate ± SE** | ***p*** | $R^2_m$ | $R^2_c$ | **Temperature (*p*)** | **Transformation** |
|---|---|---|---|---|---|---|
| ***%Sleep*** | -2.66± 5.38 | 0.622 | 0.86 | 0.90 | 0.549 | - |
| ***Frequency of sleep bouts*** | -0.13±0.26 | 0.627 | 0.54 | 0.57 | 0.054 | - |
| ***Duration of sleep bouts*** | 0.10±0.28 | 0.722 | 0.50 | 0.65 | 0.548 | Log |
| ***Interval between sleep bouts*** | 0.10±0.27 | 0.719 | 0.44* | 0.44* | 0.737 | Log |
