## Supporting Information 2 for "How variable and stress-sensitive is sleep expression in lizards?"

**Sleep-wake classifications (S, WK) based on EOG density in *Psammophilus dorsalis* lizards (n = 19).**

**Lizard ID: S01; Population origin: Rural**

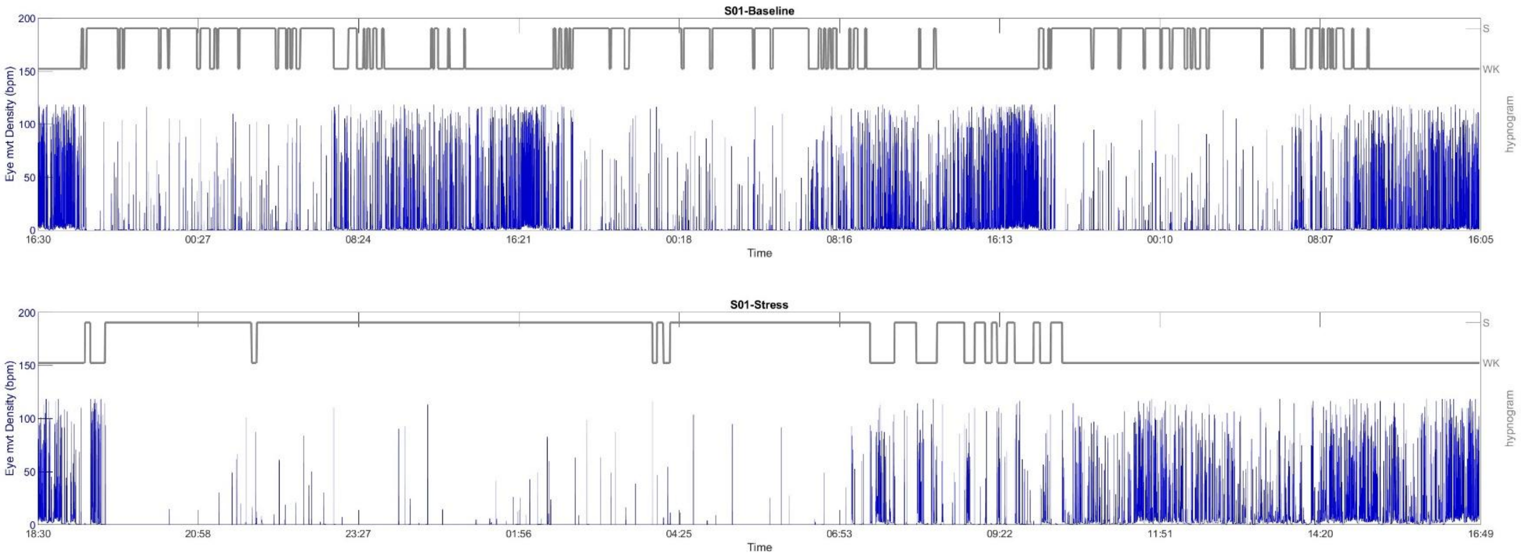

**Lizard ID: S02; Population origin: Rural**

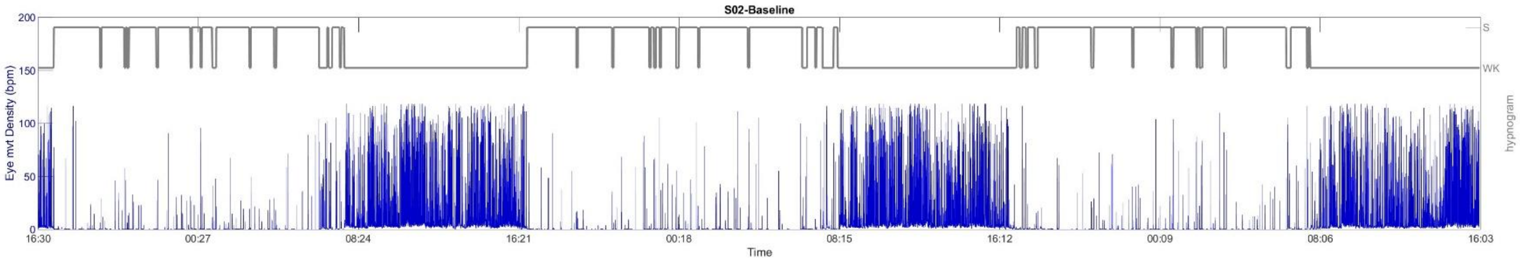


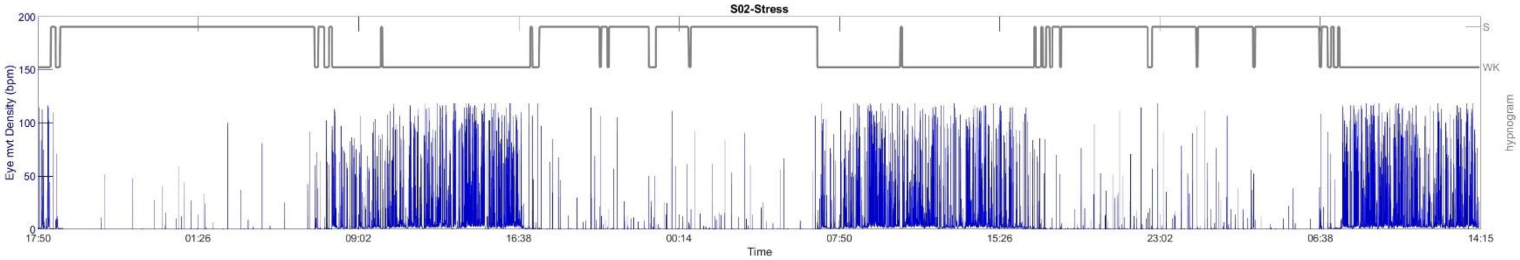

**Lizard ID: S06; Population origin: Urban**

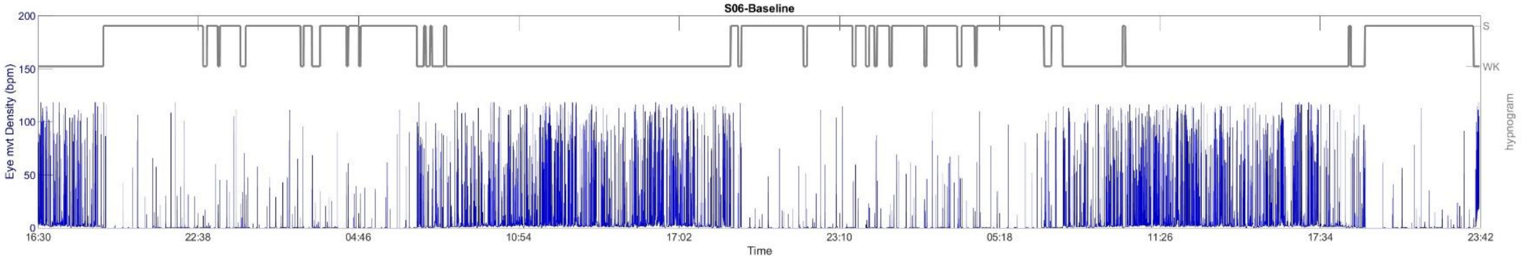

#### Lizard ID: S08; Population origin: Rural

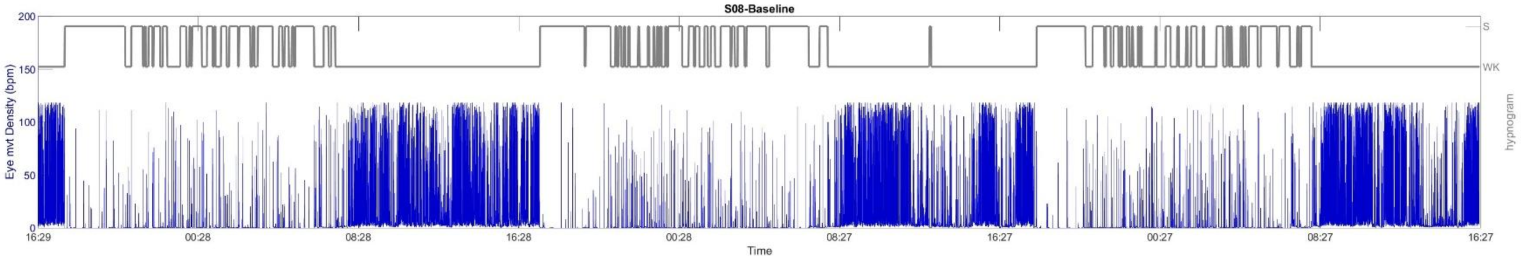

**Lizard ID: S09; Population origin: Rural**

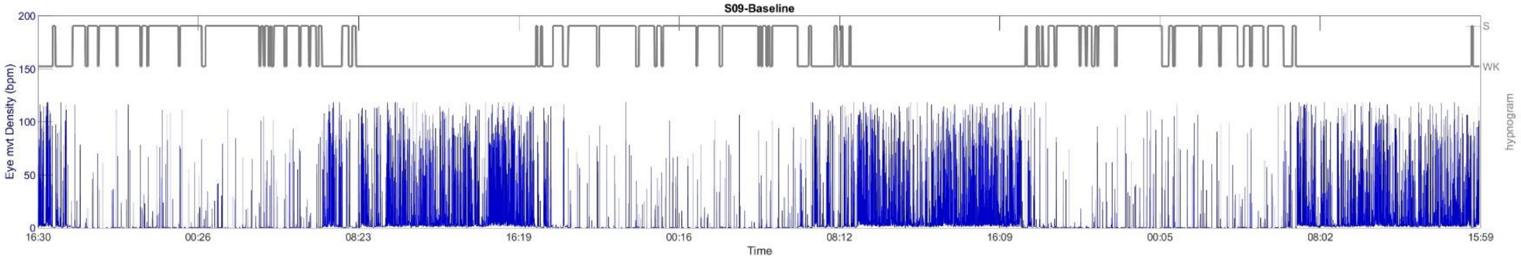


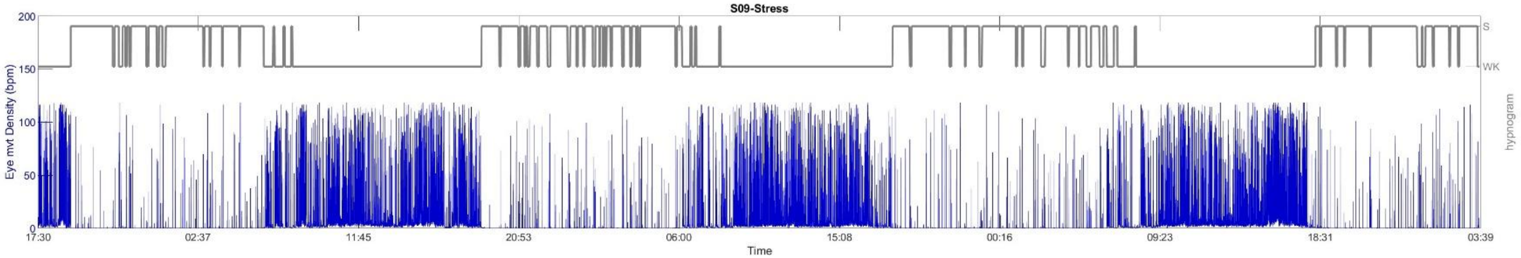

**Lizard ID: S10; Population origin: Urban**

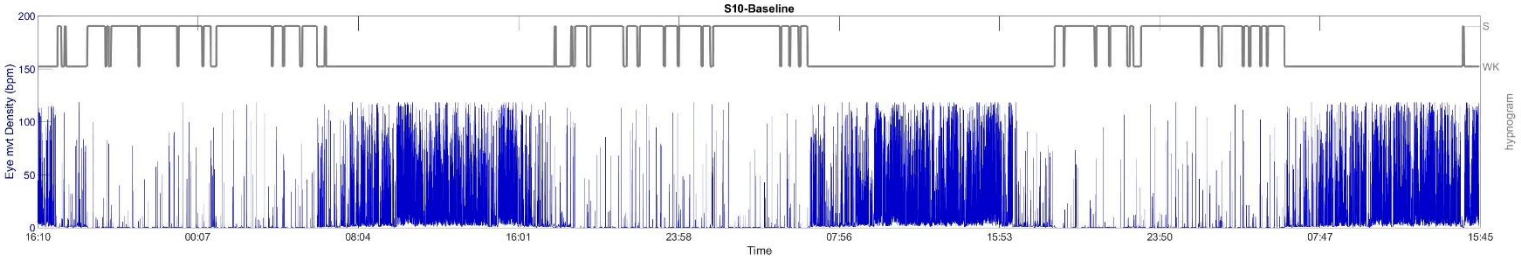


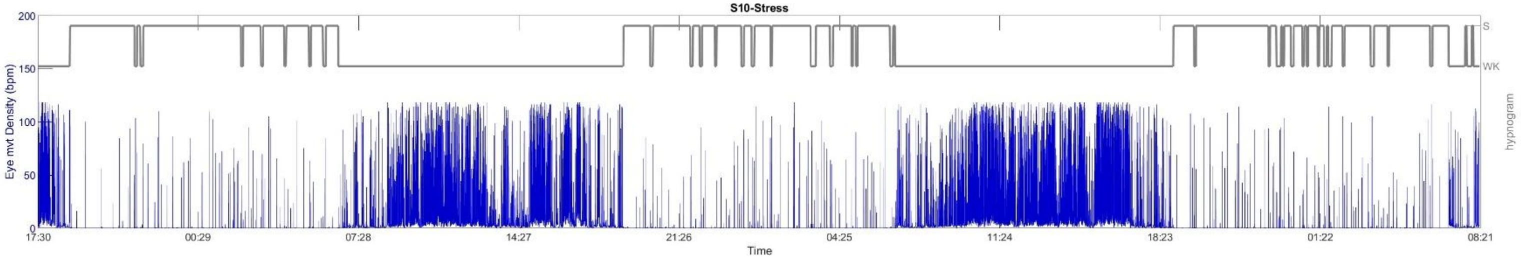

### Lizard ID: S11; Population origin: Urban

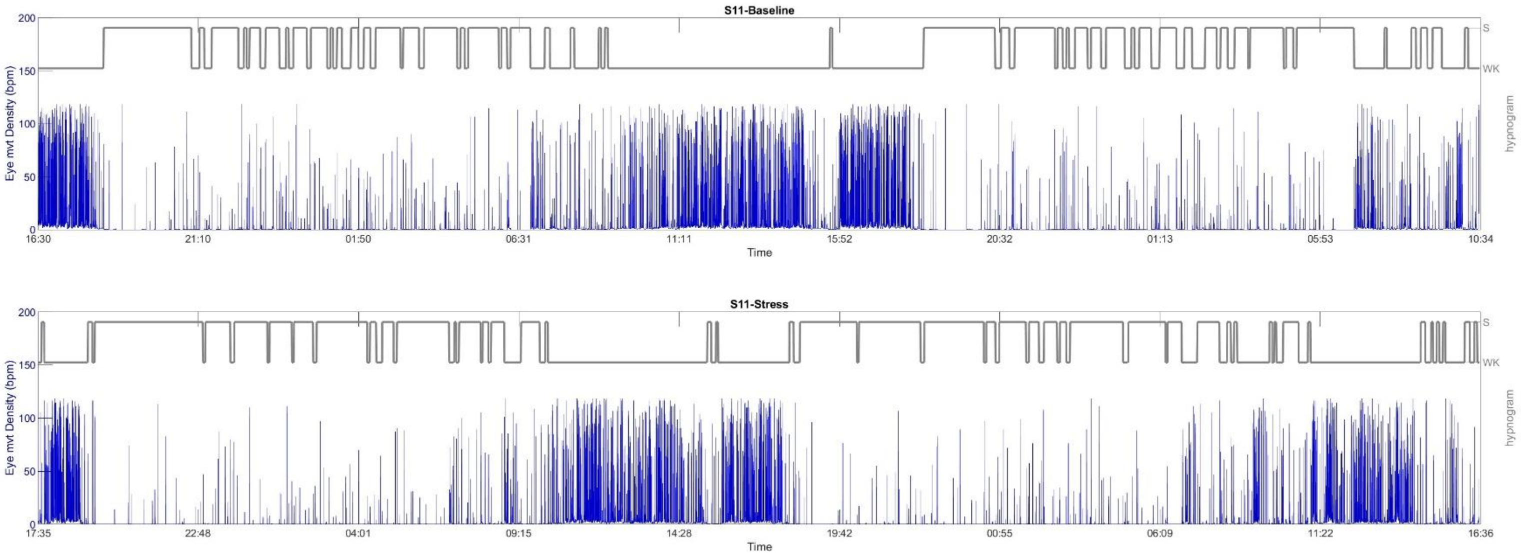

**Lizard ID: S12; Population origin: Rural**

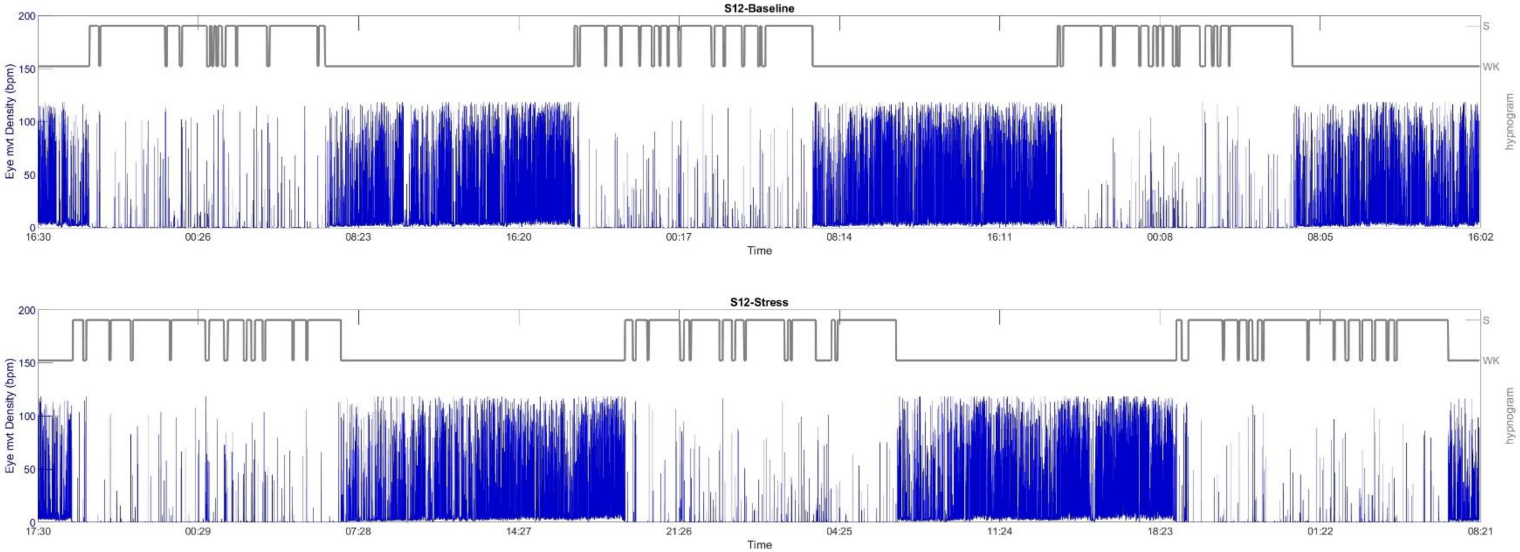

#### Lizard ID: S14; Population origin: Urban

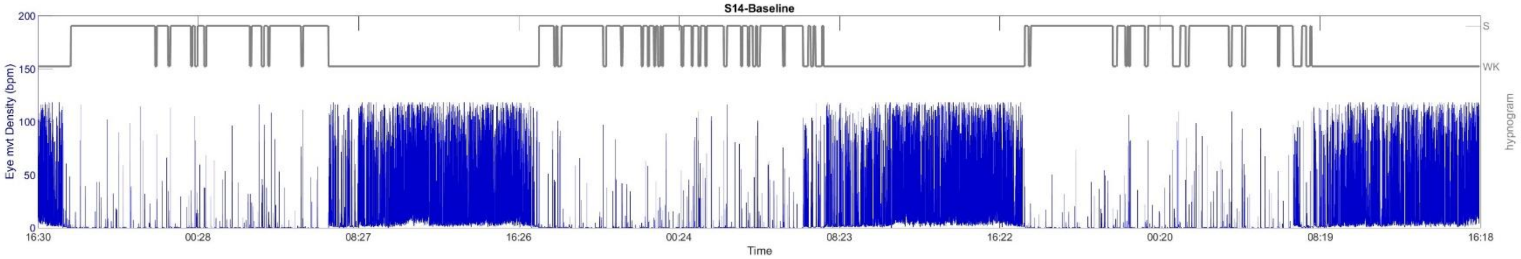


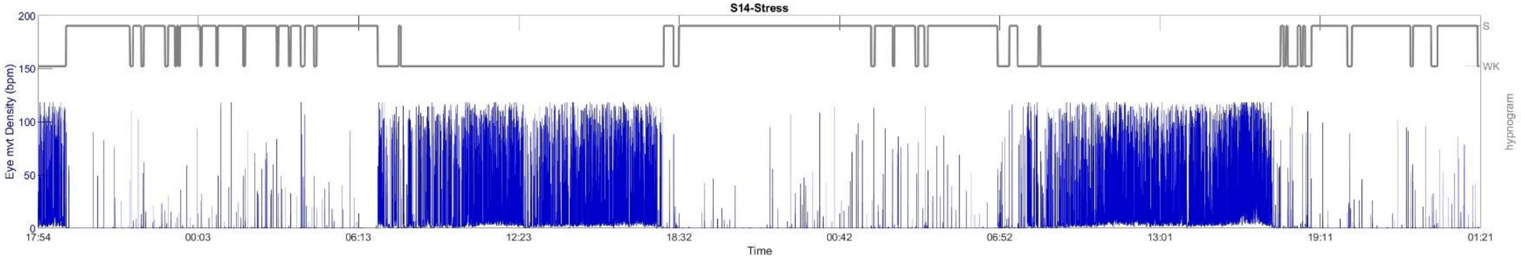

**Lizard ID: S15; Population origin: Urban**

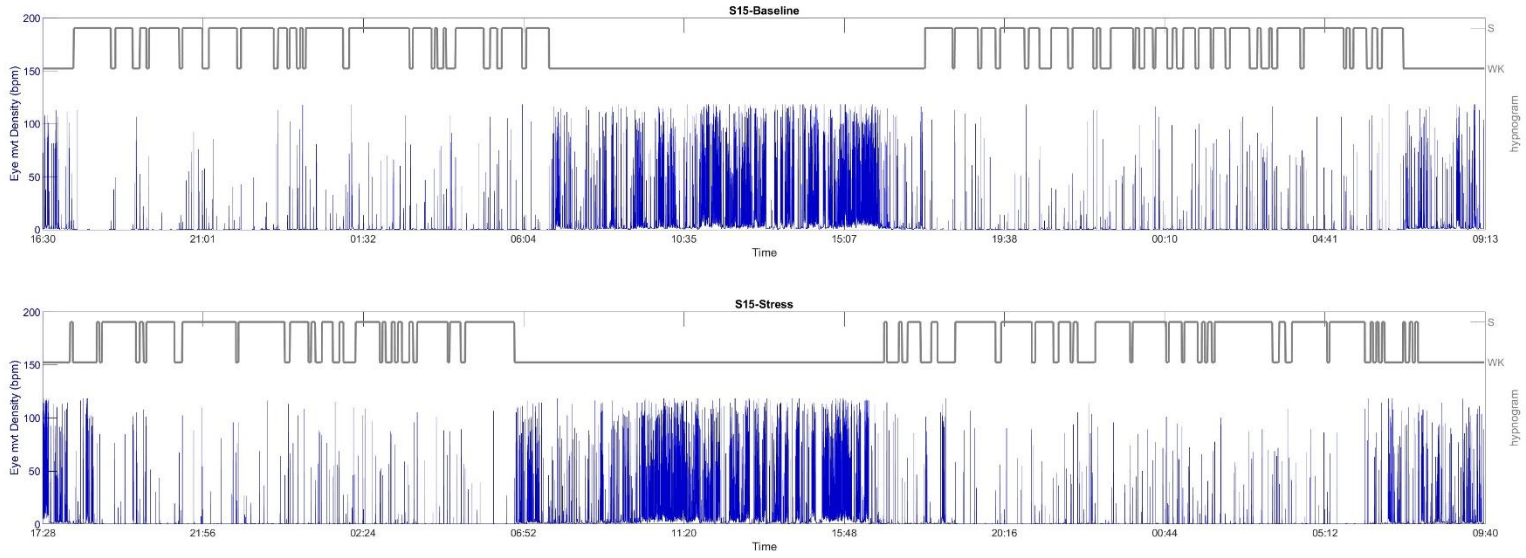

**Lizard ID: S16; Population origin: Urban**

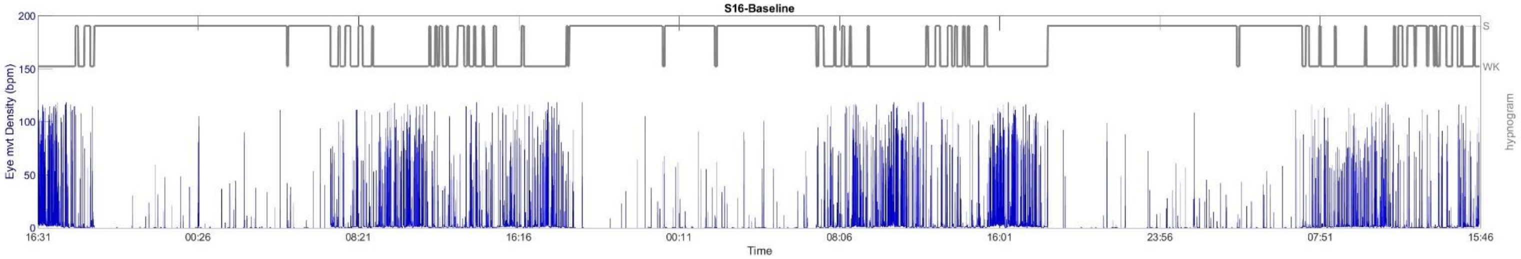


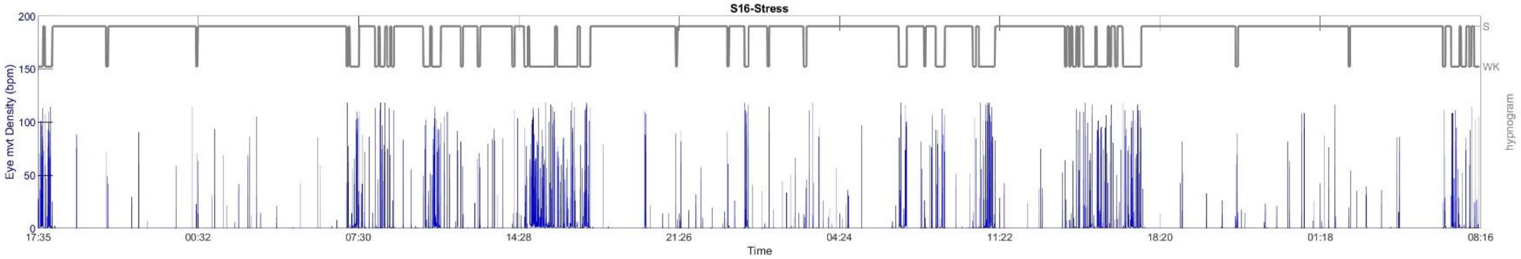

**Lizard ID: S17; Population origin: Urban**

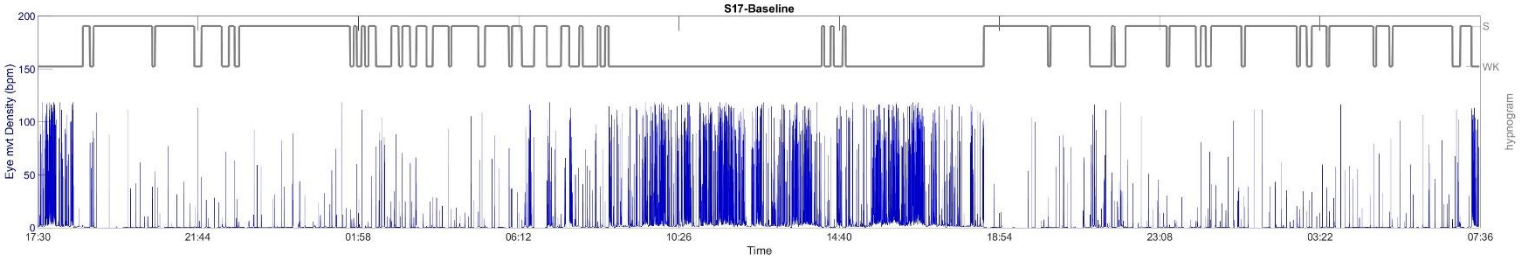


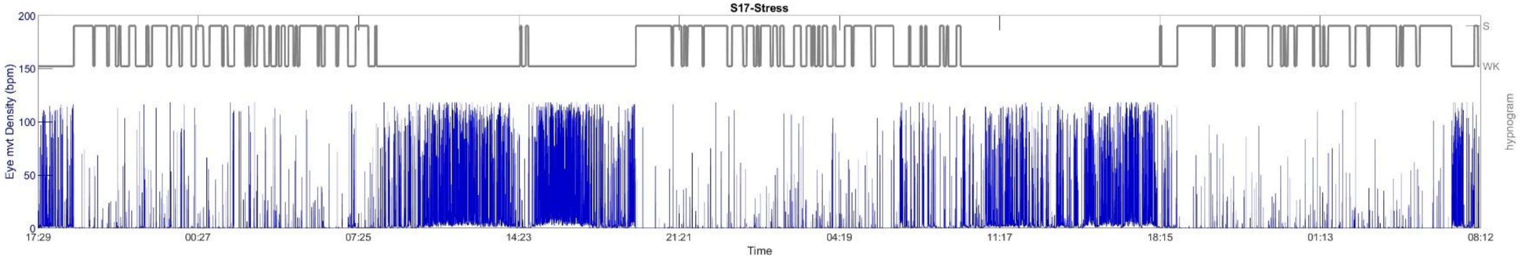

**Lizard ID: S18; Population origin: Rural**

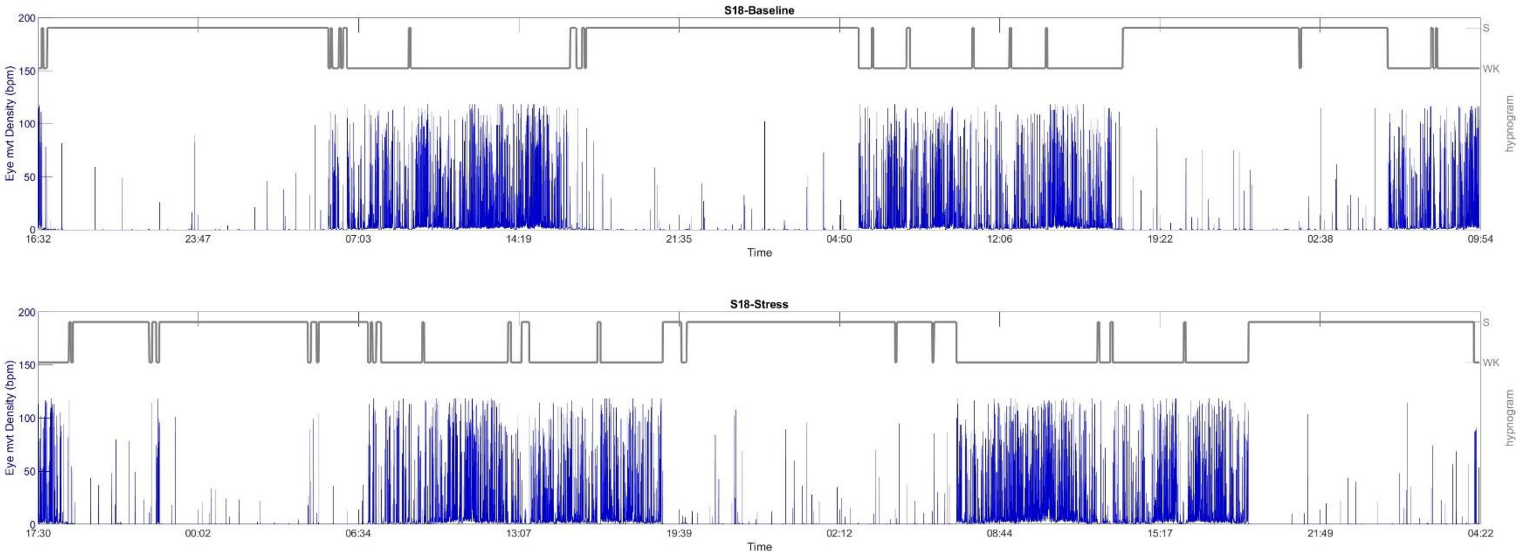

#### Lizard ID: S19; Population origin: Rural

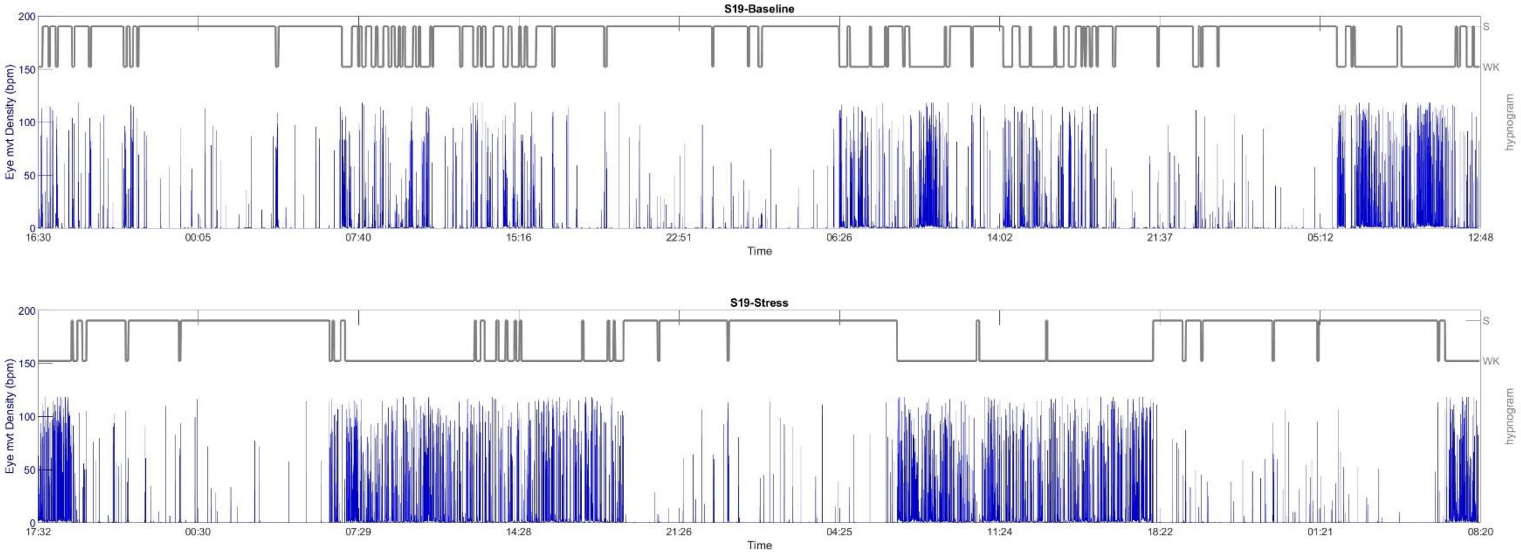

**Lizard ID: S20; Population origin: Urban**

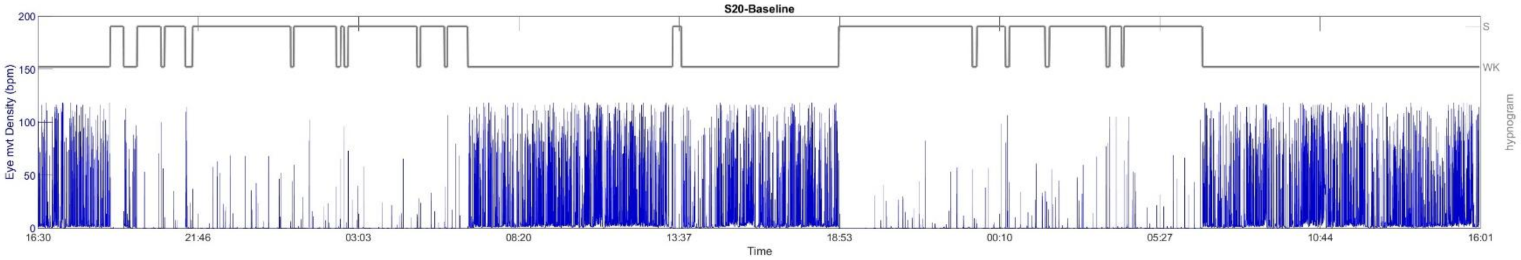

**Lizard ID: S21; Population origin: Urban**

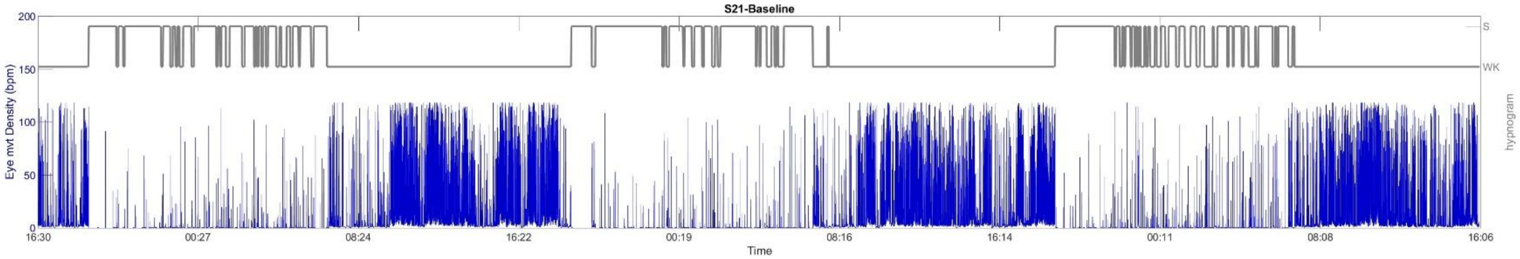


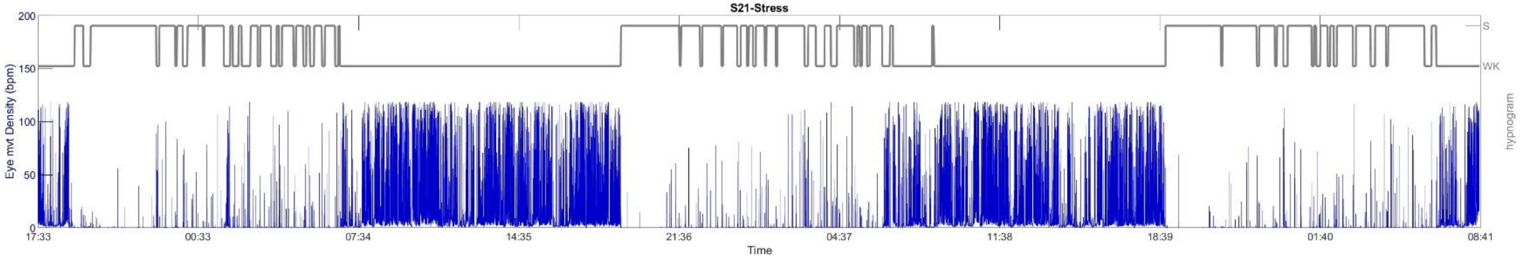

#### Lizard ID: S22; Population origin: Rural

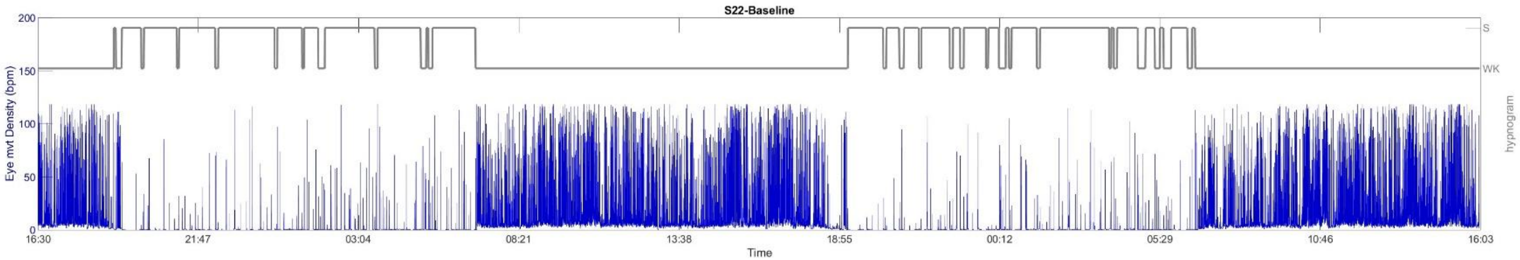

**Lizard ID: S23; Population origin: Rural**

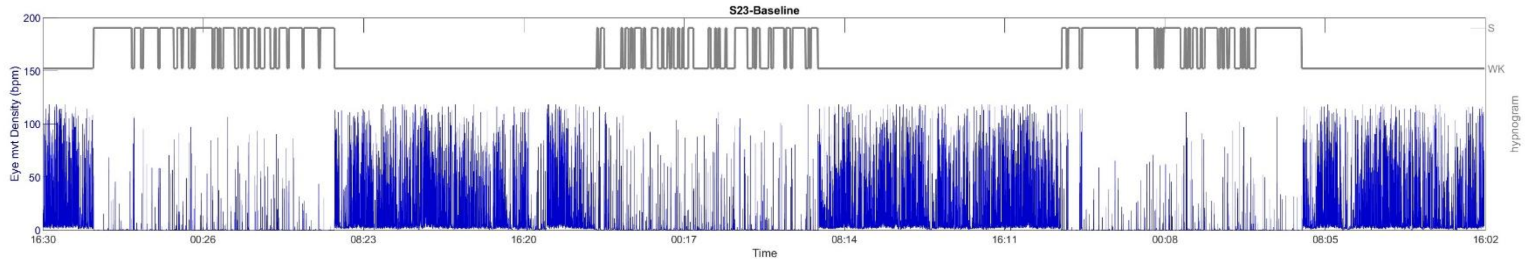

**Lizard ID: S24; Population origin: Urban**

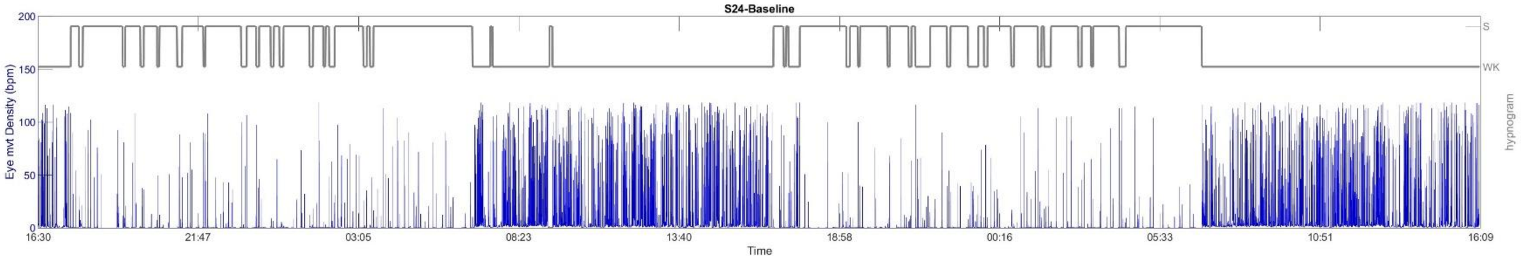
